# Emergence of tunneling nanotube-like intervesicular connections

**DOI:** 10.64898/2026.08.10.743838

**Authors:** Sungwoo Han Park, Sang Ho Lee, Chang Ho Kim, Chengying Yin, Xue Ke, Liangfei Tian, Kwanwoo Shin

**Affiliations:** Department of Chemistry and Institute of Biological Interface, Sogang University; Seoul, 04107, Republic of Korea; Zhejiang Key Laboratory of Intelligent Sensing Technology and Advanced Medical, Instrument and Key Laboratory for Biomedical Engineering of Ministry of Education, College of Biomedical Engineering & Instrument Science, Zhejiang University; Hangzhou 310027, PR China

## Abstract

Tunneling nanotubes (TNTs) are actin-supported membrane bridges that mediate long-range intercellular communication and direct transfer of signaling molecules, organelles, and pathogenic cargo, yet the physicochemical mechanisms underlying their formation and organization remain poorly understood. Here we show that TNT-like intervesicular connections emerge from minimal physicochemical interactions between actin filaments and lipid membranes. Upon Mg²⁺ exposure, actin-encapsulating vesicles spontaneously generated actin bundle-embedded lipid nanotubes (AT-LNTs) that formed stable intervesicular networks. Mg²⁺ simultaneously induced actin polymerization, filament bundling, and electrostatic recruitment of F-actin to phosphatidylcholine membranes, enabling membrane tubulation without actin-binding proteins. Systematic perturbation of membrane phase, membrane tension, Mg²⁺ concentration, ionic strength, and actin concentration revealed that AT-LNT formation occurs only within a narrow physicochemical regime where membrane deformation and actin-membrane coupling are simultaneously permissive. The resulting AT-LNTs reproduced key structural and dynamic features of cellular TNTs, including bundled organization, helical unwinding, lumenal diffusion, and spontaneous bridging between synthetic vesicles and living cells. These findings establish a minimal biophysical framework for understanding the emergence of intercellular membrane connections in living systems and provide a foundation for engineering communication between synthetic and living cell.

## Main Text

Direct physical connections between cells can form through thin membranous bridges known as tunneling nanotubes (TNTs), enabling long-range intercellular communication and direct exchange of cellular materials (*1*). These submicrometer membrane tubes can span tens of micrometers between distant cells and have been implicated in organelle transfer (*2*, *3*), signal propagation (*4*, *5*), and the intercellular spread of pathogens such as viruses (*6*), oncogenic cargo (*7*), and prions (*8*). Structurally, TNTs consist of actin filaments (F-actin) inside the lumen of the tubular membrane that provide mechanical stability and tracks for cargo transport (*9*). More broadly, actin-driven membrane tubules arise in diverse cellular processes, including filopodia (*10*, *11*), endoplasmic reticulum network (*12*), and endocytic fusion or fission intermediates (*13*), underscoring the central role of actin-membrane interactions in shaping cellular membrane architectures. Despite their biological importance, the physical mechanisms by which actin generates and stabilizes TNT-like intercellular membrane tubes remain poorly understood (*14*).

To investigate how actin drives membrane tubulation, a variety of *in vitro* reconstitution systems have been developed using lipid vesicles (*15*). Previous studies demonstrated that membrane protrusions can arise through mechanical coupling between actin assembly and membrane elasticity, either via encapsulated actin bundles (*16*, *17*) or through actin network growth on membrane surfaces (*18*, *19*). Although these studies have provided important insights into actin-driven membrane deformation, they do not reproduce a characteristic feature of TNTs, the spontaneous formation of long intervesicular membrane tubes supported by internal actin bundles that directly connect separate membrane compartments (*9*, *14*). Consequently, existing *in vitro* systems have yet to recapitulate the spontaneous formation of TNT-like membrane connections between separate membrane compartments.

Here we show that TNT-like *inter*vesicular membrane connections can emerge from minimal physicochemical interactions between actin filaments and lipid membranes. Upon Mg²⁺ exposure, lipid vesicles encapsulating actin monomers spontaneously generate actin bundle-embedded lipid nanotubes (AT-LNTs) that connect neighboring vesicles. Mg²⁺ simultaneously promotes actin polymerization, filament bundling, and electrostatic recruitment of F-actin to phosphatidylcholine (PC) membranes, thereby enabling membrane tubulation without actin-binding proteins. By systematically varying membrane phase, membrane tension, Mg²⁺ concentration, ionic strength, and actin concentration, we identify the physicochemical conditions governing nanotube formation and stabilization. The resulting AT-LNTs form TNT-like bundled conformations, undergo dynamic remodeling, and permit lumenal diffusion of solutes. Notably, AT-LNTs also extended toward living cells and formed stable vesicle-cell connections, accompanied by cellular pulling and localized actin recruitment at the docking sites. This spontaneous coupling suggests that our identified physicochemical principles may capture fundamental features of TNT emergence. Our findings establish a minimal, tunable platform for dissecting the biophysical basis of intercellular connectivity and for engineering direct communication between synthetic and living cells.

### Mg²⁺-mediated actin-membrane interactions generate TNT-like intervesicular nanotubes

To investigate whether actin-membrane interactions alone can generate TNT-like membrane connections, we encapsulated soluble actin monomers (G-actin) at 1 μM, near the critical concentration for filament elongation (*20*, *21*), in lipid vesicles composed of liquid-disordered (L_d_) PC (dioleoylphosphatidylcholine, DOPC) membrane embedded with Mg²⁺ ionophores (5 mol%) **(Fig. 1A, a)**. A small aliquot of these actin-containing vesicles (hereafter actin-vesicles) was transferred into external buffer containing 20 mM Mg²⁺ **(fig. S1A)**, consistent with the reported concentration range required for actin bundling (*22*) and effective charge modulation of phosphatidylcholine membranes (*23*, *24*), thereby enabling actin-membrane association (*25*).

**Figure 1.**
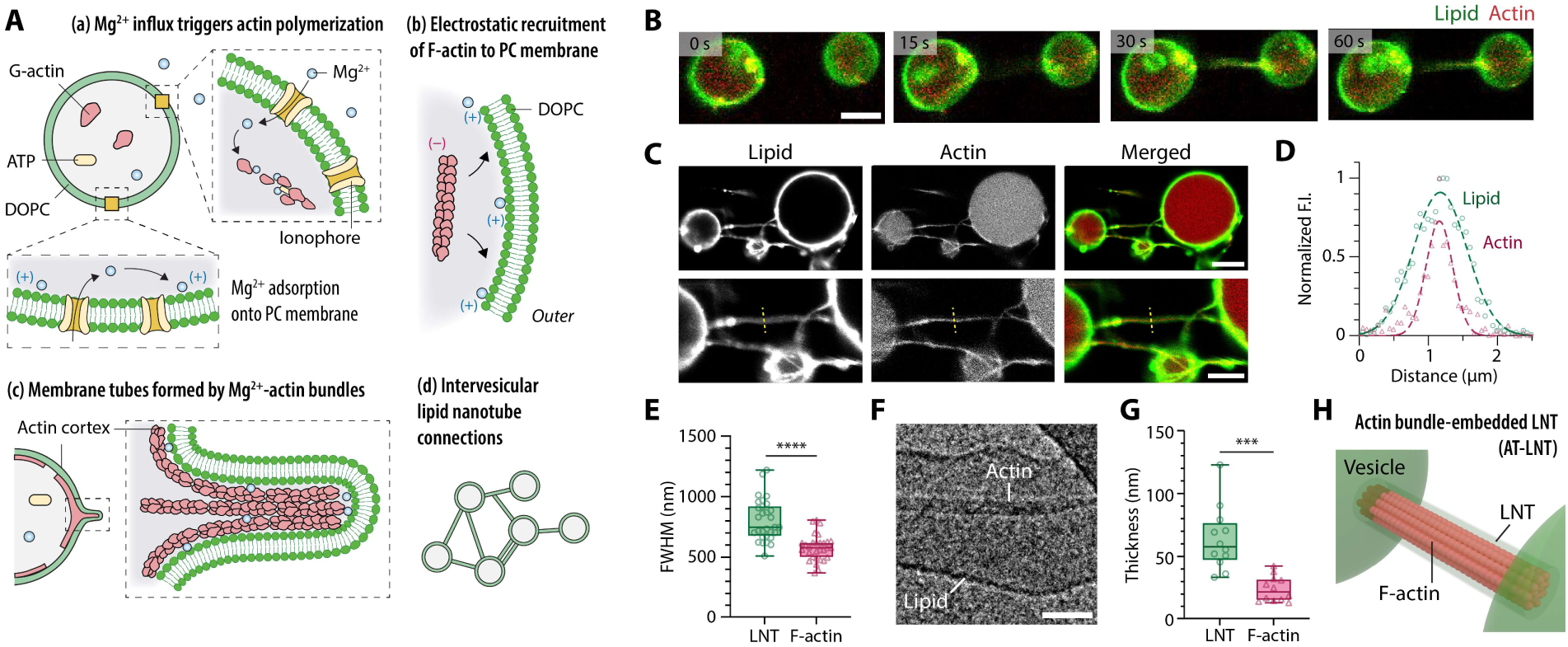
Mg^2+^-mediated formation and structural characterization of actin bundle-embedded lipid nanotubes (AT-LNTs). **(A)** Schematic of Mg²⁺-mediated AT-LNT formation. (a) Mg²⁺ permeates into actin-encapsulating vesicles, adsorbs onto the inner leaflet of the PC membrane, and triggers polymerization. (b) The PC membrane recruits negatively charged F-actin via Mg²⁺-mediated electrostatic attraction. (c) Mg²⁺-crosslinked actin bundles push the membrane outward, generating membrane tubes. (d) Tubules elongate to form intervesicular connections. **(B)** Representative time-lapse images of intervesicular AT-LNT formation. Green, lipid; red, actin. Scale bar, 5 μm. See also movie S2. **(C)** Representative images of AT-LNT networks. Green, lipid; red, actin. Scale bars, 10 μm (top) and 5 μm (bottom). **(D)** Normalized fluorescence intensity (F.I.) profiles of lipid (green) and actin (red) along the yellow dashed line in (C). See Fig. S7B for additional profiles. **(E)** FWHM comparison of lipid and actin signals measured at perpendicular cross-sections. Mean ± SD; *n =* 30; \*\*\*\**p* < 0.0001, two-tailed t-test. **(F)** Representative cryo-EM image of an AT-LNT. Scale bar, 50 nm. **(G)** Thickness of LNTs and internal actin bundles measured by cryo-EM. Mean ± SD; *n =* 12; \*\*\**p* < 0.001, two-tailed t-test. **(H)** Proposed structural model of AT-LNTs, showing an actin bundle embedded within the LNT lumen.

Mg²⁺ rapidly permeated into the vesicle lumen through the embedded ionophores **(Fig. 1A, a; fig. S1, B and C)**, where it induced actin polymerization (*26*) and promoted crosslinking of the resulting filaments into bundles. Simultaneously, Mg²⁺ adsorption onto PC membranes induced electrostatic attraction between negatively charged F-actin and the membrane surface, leading to accumulation of cortical F-actin **(Fig. 1A, b and c and fig. S2, A-C**

We confirmed the electrostatic nature of this interaction using negatively charged dioleoylphosphatidylglycerol (DOPG) membranes **(fig. S2D)**. Under identical 20 mM Mg²⁺, F-actin was expelled from DOPG membranes, resulting in actin bundles condensed toward the vesicle center rather than association with the membrane **(fig. S2E)**. Similarly, in phase-separated vesicles containing both *Ld* PG and liquid-ordered (*Lo*) PC domains, actin selectively localized to the PC-rich domains **(fig. S2, F and G)**. These results established Mg²⁺-mediated electrostatic attraction as the primary determinant of actin-membrane association in this system.

Under standard conditions (DOPC membranes, 1 μM G-actin, 20 mM Mg²⁺), abundant outward tubular structures spontaneously emerged from actin-vesicles **(Fig. 1, A-C)**. These structures were identified as lipid nanotubes (LNTs) based on their visibility under differential interference contrast (DIC) microscopy **(fig. S3)** and their characteristic thermal undulation (*27*) and retraction (*28*) under strong fluorescence illumination **(movie S1)**. Notably, many LNTs elongated toward neighboring vesicles and established stable intervesicular connections resembling tunneling nanotubes (TNTs) **(Fig. 1B and movie S2)**. Docking onto preexisting LNTs additionally generated branched configurations **(fig. S4 and movie S3)**. Additional LNTs frequently emerged near docking or growth sites, generating bundled and branched LNT networks **(fig. S5 and movies S4-S6)**. In some cases, vesicles initially enclosed within larger vesicles escaped and formed new LNT connections with external structures **(fig. S6 and movie S7)**. Following docking, LNT connections underwent dynamic rearrangements including pulling, detachment, and reconnection, eventually forming stable intervesicular networks that persisted for several days **(movie S8)**.

Fluorescence imaging revealed strong colocalization of lipid and actin signals along intervesicular LNTs **(Fig. 1C)**. Cross-sectional intensity profiles showed overlapping lipid and actin signals **(Fig. 1D and fig. S7, A and B)**. Notably, the full width at half maximum (FWHM) of the lipid signals (805 ± 171 nm) remained significantly greater than that of the actin signals (579 ± 107 nm) **(Fig. 1E)**, indicating that actin filaments occupy the LNT lumen rather than the membrane surface. We further examined FWHM pairs at 11 positions along a single intervesicular LNT **(fig. S7C)**. While the LNT (828 ± 101 nm, ± 12%) exhibited more homogeneous thickness than the internal actin (414 ± 167 nm, ± 40%), significant correlation existed between paired measurements **(fig. S7D)**, suggesting structural coupling between internal actin organization and nanotube morphology. Finally, three-dimensional reconstruction confirmed that actin filaments are enclosed within the LNT membrane and that the LNTs are positioned above the substrate **(fig. S7E and movie S9)**.

To directly resolve the internal LNT architecture, we performed cryo-electron microscopy (cryo-EM) **(Fig. 1F)**. The LNTs contained bundled actin filaments aligned along the lumen, reproducing a defining structural feature of cellular TNTs (*9*). At growth or docking sites, aligned actin bundles or fragmented filaments were frequently oriented along the nanotube axis **(fig. S8)**. The nanotube diameter ranged 33-123 nm, with a mean diameter 64 ± 25 nm, whereas internal actin bundle diameters ranged 13-42 nm, with a mean diameter 24 ± 10 nm **(Fig. 1G)**. Based on these dimensions, the estimated persistence lengths of the internal actin bundles (30-360 µm; **Supplementary Note 1**) indicate sufficient mechanical rigidity to support the long-range intervesicular connections observed by confocal microscopy. We therefore designate these structures as actin bundle-embedded lipid nanotubes (AT-LNTs) **(Fig. 1H)**.

### Membrane energetics governs AT-LNT formation

Vesicle-tube systems follow classical membrane mechanics, where the force required to form a membrane nanotube is given by 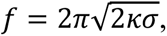 with κ the bending modulus and *σ* the membrane tension (*29*, *30*). We first examined the effect of membrane rigidity by varying lipid phase composition. *Lo* phase vesicles composed of distearoylphosphatidylcholine (DSPC) and cholesterol (70:30 mol%) (*31*), which retain the same phosphocholine group but exhibit substantially higher bending rigidity than DOPC membranes (*32*, *33*) **(Fig. 2A)**, formed membrane-associated actin cortices but failed to generate AT-LNTs **(Fig. 2, B and C)**. This indicates that actin-membrane association alone is insufficient for nanotube formation when membrane deformation energy is prohibitively high.

**Figure 2.**
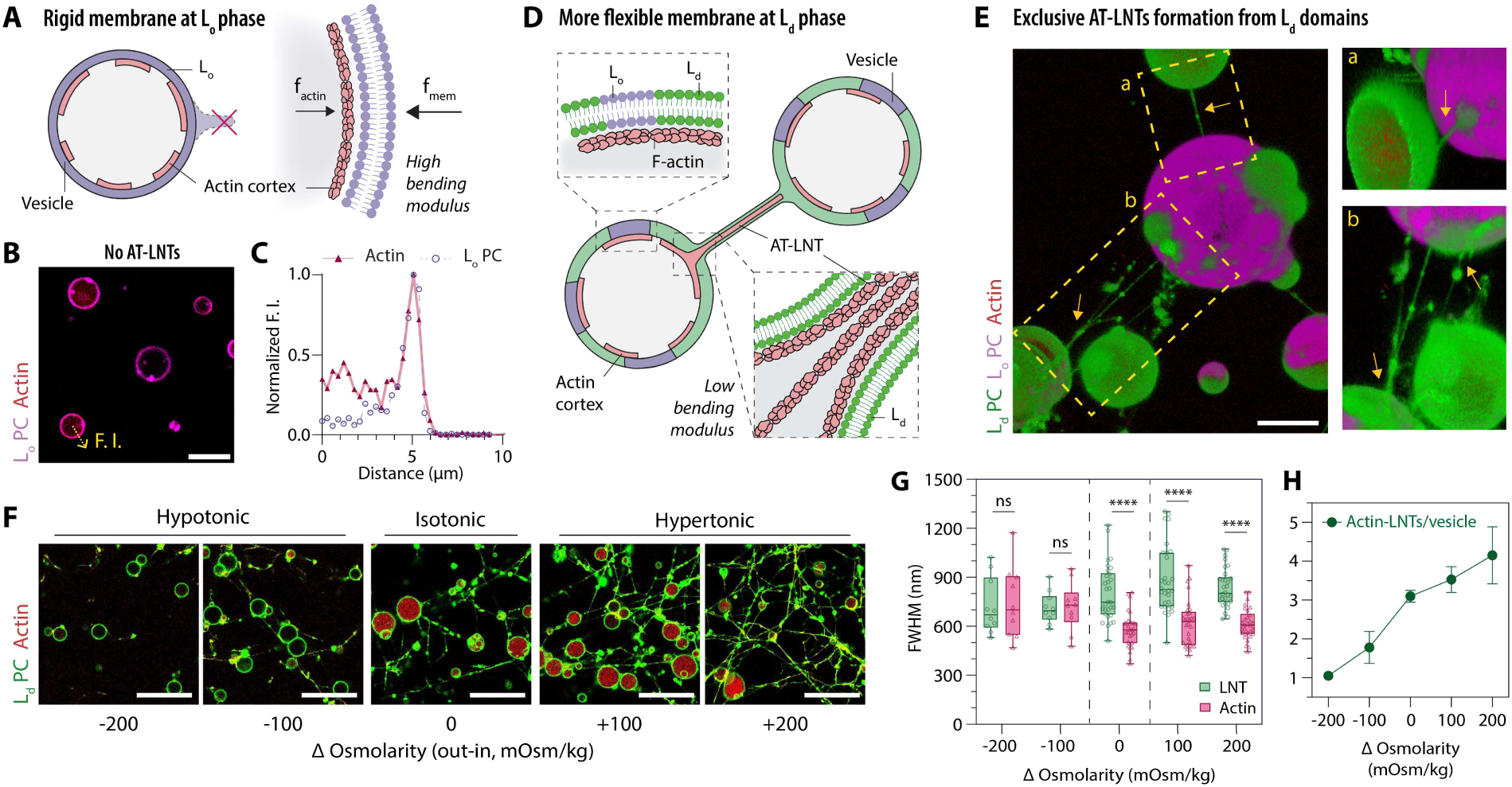
AT-LNT formation under different membrane energetic conditions. **(A)** Schematics showing that AT-LNTs cannot form from actin-vesicle with an L_o_ phase PC membrane due to its high bending modulus (n ↑). **(B)** Representative confocal fluorescence image of actin-vesicles with L_o_ PC membrane in 20 mM Mg^2+^. No AT-LNTs are observed. Magenta, L_o_ membrane; red, actin. Scale bar, 20 μm. **(C)** Normalized fluorescence intensity profiles of lipid and actin along the yellow dashed arrow in (B). The actin peak colocalizes with the L_o_ PC membrane. **(D)** Schematics showing exclusive AT-LNT formation from the L_d_ domain with lower bending modulus (n ↓) of the phase-separated vesicles. **(E)** 3D reconstruction of AT-LNTs formed from actin-vesicles with phase-separated PC membrane in 20 mM Mg^2+^, generated from Z-stacks of confocal fluorescence microscopy images. AT-LNTs (yellow arrows) are composed of L_d_ membrane and are exclusively formed from or docked onto L_d_ domains. Green, L_d_ membrane; magenta, L_o_ membrane; red, actin. Scale bar, 10 μm. **(F)** Intervesicular AT-LNT network formed in 20 mM Mg^2+^ buffer under different osmotic conditions. Scale bars, 50 μm. **(G)** Cross-sectional FWHMs of lipid and actin signals from AT-LNTs under different osmotic conditions. Data represent mean ± SD; *n* = 10 per hypotonic condition and *n* = 30 per isotonic and hypertonic condition; \*\*\*\**p* < 0.0001, two-tailed t-test. **(H)** Average number of AT-LNTs per vesicle under different osmotic conditions. Data represent mean ± SD; *n* = 3 images (387.5 μm × 387.5 μm) per condition.

To directly compare membrane phase behavior within the same vesicle, we generated phase-separated vesicles with coexisting *Ld* and *Lo* domains of PC membrane **(Fig. 2D)**. AT-LNTs were exclusively composed of *Ld* membrane, with both nanotube growth and docking sites spatially restricted to the *Ld* domains **(Fig. 2E)**. Because *Ld* membranes possess substantially lower bending rigidity than *Lo* membranes, these results establish membrane deformability as a key energetic requirement for AT-LNT formation.

Having established membrane phase as a permissive factor for AT-LNT formation, we next examined whether modulating the overall osmotic environment, and thus the resulting membrane tension could further promote tubule emergence by altering the mechanical context. Under hypoosmotic conditions (*C_osm_* = *C_out_* − *C_in_* = −100, −200 mOsm/kg) across the vesicles, which increase membrane tension, actin bundles were frequently observed outside vesicles as leaked actin **(Fig. 2F and fig. S9)**. Fragmented lipid signals showed only sparse colocalization with external actin bundles, without significant differences in FWHM thickness at colocalization sites **(Fig. 2G)**, indicating that membrane tubes containing internal actin bundles were no longer formed. These observations suggest that elevated membrane tension suppresses stable membrane tubulation despite continued actin polymerization and bundling.

In contrast, hyperosmotic conditions (*C_osm_* = +100, +200 mOsm/kg), which reduce membrane tension, produce significantly more AT-LNTs **(Fig. 2H)** and more extensive intervesicular networks **(Fig. 2F)**. Consistent with classical membrane mechanics, where nanotube radius follows 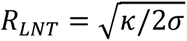, nanotube diameters increased slightly under reduced membrane tension (885 ± 206 nm and 835 ± 118 nm for +100 and +200 mOsm/kg, respectively) compared to isotonic conditions (805 ± 171 nm), although these differences were not statistically significant **(Fig. 2G)**. The limited diameter increase likely reflects energetic dispersion across the multiple AT-LNTs generated from a vesicle under low-tension conditions (*34*, *35*).

Together, these results demonstrate that AT-LNT formation is governed by the membrane energetics of vesicle-tethered nanotubes and that both membrane rigidity and tension define the physical regime permissive for TNT-like intervesicular nanotube formation.

### Physicochemical regime for AT-LNT formation

To dissect the specific contribution of Mg²⁺ from that of actin, we examined Mg²⁺-treated DOPC vesicles lacking actin. Upon exposure to 20 mM Mg²⁺, thin LNTs (FWHM 517 ± 150 nm) spontaneously formed and exhibited characteristic thermal undulation **(fig. S10A and movie S10)**. Although most nanotubes gradually disappeared through retraction or adhesion to substrates **(fig. S10B)**, a subset that established intervesicular connections became stabilized. Vesicle aggregation and hemifusion were also frequently observed **(fig. S10A)**, indicating that Mg²⁺ alone can induce both LNT formation and intermembrane adhesion (*36–38*), thereby providing a possible physical basis for nanotube docking onto neighboring vesicle surfaces. Henceforth, we designate these actin-free, thin, and thermally fluctuating nanotubes as Mg²⁺-induced lipid nanotubes (Mg-LNTs).

We next varied Mg²⁺ concentration for actin-vesicles containing 1 μM actin. In the absence of Mg²⁺, no nanotube formation was observed **(fig. S11A)**. At 5 mM Mg²⁺, where Mg²⁺-induced charge modulation of phosphatidylcholine membranes is minimal (*23*), only thin Mg-LNTs formed (FWHM 552 ± 98 nm) without detectable actin colocalization **(fig. S11, A and B; movie S11)**. In contrast, at 10-50 mM Mg²⁺, where Mg^2+^-driven membrane charge modulation enables actin-membrane association (*25*), thicker AT-LNTs (FWHM 844 ± 218 nm at 10 mM Mg^2+^) containing internal actin filaments formed with morphologies identical to those observed under standard conditions **(fig. S11A)**. At 100 mM Mg²⁺, unstable AT-LNTs and Mg-LNTs exhibiting repeated reconnection events were observed **(fig. S11, A and C; movie S12)**, suggesting that excessive Mg²⁺ impairs electrostatic actin-membrane association and persistent AT-LNT organization via charge-screening effect (*39*). This morphological transition spanning Mg-LNTs at ∼5 mM Mg^2+^, AT-LNTs at 10-50 mM Mg^2+^, and their coexistence at 100 mM Mg^2+^ indicates that Mg^2+^-mediated actin-membrane association is the key determinant for AT-LNT formation **(fig. S11D)**.

To directly modulate electrostatic interactions through charge-screening effects (*39*), we increased ionic strength by adding NaCl (**Fig. 3A**). AT-LNT formation persisted up to 82.5 mM ionic strength, whereas at ≥105 mM, numerous Mg-LNTs lacking internal actin signal were observed (**Fig. 3, B and C**; **fig. S12A**; **movie S13**). Nanotube thickness progressively decreased with increasing ionic strength, from 795 ± 120 nm under Na⁺-free conditions to 607 ± 91 nm at physiological 150 mM ionic strength (**Fig. 3B**). These results indicate that electrostatic screening weakens actin-membrane coupling and progressively shifts the system from actin-supported AT-LNTs toward thinner Mg-LNT structures. Reduced actin-membrane interactions additionally enabled real-time observation of Mg-LNT formation (**fig. S12B** and **movie S14**).

**Figure 3.**
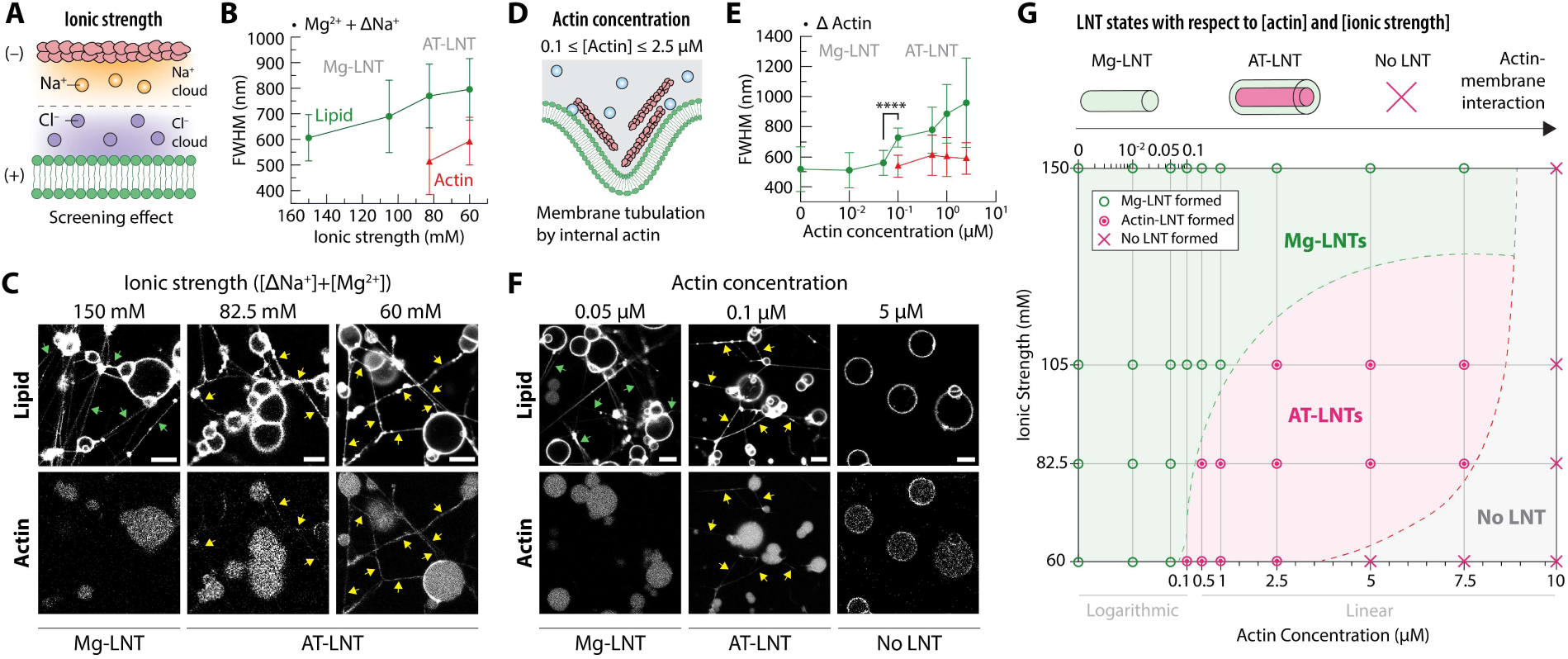
Ionic strength and actin concentration control AT-LNT formation. **(A)** Schematics showing electrostatic screening between actin and membrane at elevated NaCl concentrations. **(B)** Cross-sectional FWHMs of lipid and actin signals from nanotubes as a function of ionic strength (*n* = 10 tubules per condition; data represent mean ± SD). **(C)** Representative confocal fluorescence images of Mg-LNTs (green arrows) and AT-LNTs (yellow arrows) formed from actin-vesicles (1 μM actin) at varying ionic strength (20 mM Mg²⁺). Scale bars, 10 μm. **(D)** Schematics of AT-LNT formation at an optimal actin concentration. **(E)** Cross-sectional FWHMs of lipid and actin signals from nanotubes as a function of actin concentration (*n* = 10 tubules per condition; data represent mean ± SD; \*\*\*\**p* < 0.0001). **(F)** Representative confocal fluorescence images of Mg-LNTs (green arrows) and AT-LNTs (yellow arrows) formed from actin-vesicles with varying actin concentration (Na^+^-free 20 mM Mg^2+^ external buffer). Scale bars, 10 μm. **(G)** Phase diagram of LNT morphologies as a function of [Actin] and [Ionic strength] with regions corresponding to Mg-LNTs (green circles), AT-LNTs (red circles), and no LNT formation (red crosses). Boundaries between regions are drawn arbitrarily between points of different morphologies.

We next varied actin concentration inside vesicles (**Fig. 3D** and **fig. S13A**). At ≤ 0.05 μM actin, only thin Mg-LNTs formed (FWHMs 510-561 nm) without detectable actin colocalization (**Fig. 3, E and F**; **fig. S13B**; **movie S15**). In the range of 0.1-2.5 μM, thicker AT-LNTs (FWHMs 727-960 nm) containing internal actin bundles were consistently observed (**Fig. 3, E and F**; **fig. S13B**). At ≥ 5 μM actin, nanotube formation was completely suppressed, and instead thick actin cortices developed along the vesicle membrane (**Fig. 3F** and **fig. S13B**), suggesting that excessive actin accumulation mechanically stabilizes the membrane and increases the energetic barrier for membrane tubulation (*40*, *41*).

We then constructed a phase diagram summarizing nanotube morphologies as a function of ionic strength and actin concentration **(Fig. 3G)**. Mg-LNTs were observed under weak actin-membrane interactions, whereas no nanotube formation occurred under strong interactions. AT-LNTs formed exclusively within an intermediate regime, where actin-membrane coupling is sufficient to drive tubulation without suppressing membrane deformation.

### Bundled architecture and unwinding dynamics

In addition to the AT-LNT bundling observed during nanotube growth, the broad distribution of AT-LNT thickness (FWHMs 500-1302 nm; **fig. S14**) suggests that many intervesicular connections consist of multiple individual AT-LNTs (iAT-LNTs) rather than single nanotube structures **(Fig. 4A)**. Such bundled organization is a unique structural feature of cellular tunneling nanotubes (TNTs), which are frequently composed of bundled individual TNTs (iTNTs) (*9*, *42*, *43*).

**Figure 4.**
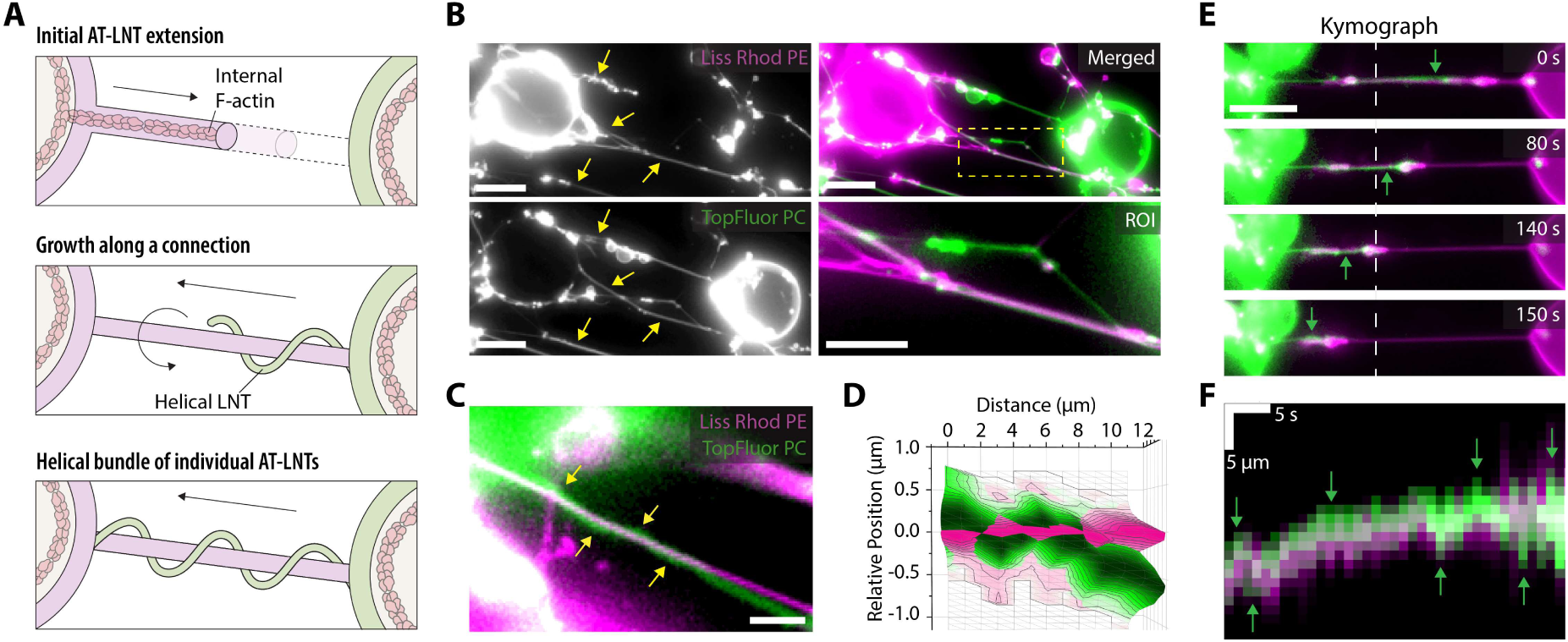
Bundled configuration and helical unwinding of AT-LNTs. Two actin-vesicle samples labeled with different lipid dyes (TopFluor^TM^ PC, green; 18:1 Liss Rhod PE, magenta) were mixed, introduced into 20 mM Mg^2+^ buffer, and imaged by fluorescence microscopy. **(A)** Schematic of helical AT-LNT bundle formation. See fig. S5B and movie S5. **(B)** Representative fluorescence images of AT-LNT bundles formed between two vesicles labeled with different lipid dyes. Individual AT-LNTs (iAT-LNTs) labeled with different lipid dyes are aligned along the connections (yellow arrows). Scale bar, 10 μm. **(C)** Representative image of an AT-LNT bundle in a helical configuration. Scale bar, 2 μm. **(D)** 3D contour plot of fluorescence intensity profiles along the helical AT-LNT bundle shown in (C). Relative position of the green AT-LNT is represented with respect to the magenta AT-LNT. **(E and F)** Unwinding process of an AT-LNT bundle triggered by photodamage to the TopFluor^TM^ PC-labeled membrane. The green AT-LNT alternated its position relative to the magenta AT-LNT over time (green arrows). **(E)** Representative time-lapse images. Times indicate duration after initiating photodamage. Scale bar, 10 μm. See also movie S16. **(F)** Kymograph measured along the white dashed line in (E). The green AT-LNT alternates its position (green arrows) relative to magenta AT-LNT over time.

To directly visualize individual nanotubes within bundled structures, we mixed actin-vesicles labeled with different lipid fluorophores and induced AT-LNT formation. Most intervesicular connections displayed alignment of both fluorescent signals along the nanotube axis **(Fig. 4B)**, demonstrating that the connections are bundles of multiple independently formed nanotubes. In some cases, individual iAT-LNTs exhibited helical organization, with one nanotube wrapping around another straight nanotube **(Fig. 4, C and D)**, producing higher-order conformations reminiscent of helical TNT bundles (*43*).

Confirming the resemblance of AT-LNTs to cellular TNTs, we observed that bundled AT-LNTs also reproduced the *unwinding dynamics* reported during the transition from bundled TNTs to single-strand TNTs in living cells (*43*). Upon inducing localized photodamage to one fluorescently labeled membrane within a bundled AT-LNT structure, the bundle underwent progressive unwinding, in which the photodamaged iAT-LNT gradually retracted toward its source vesicle while sliding along the other iAT-LNT **(Fig. 4E and movie S16)**. Kymograph analysis revealed alternating positional shifts of the photodamaged iAT-LNT over time, confirming that unwinding proceeds through helical deformation dynamics **(Fig. 4F)**. Together, these results demonstrate that higher-order TNT-like organization and dynamic remodeling can emerge from minimal actin-membrane interactions alone.

### Intervesicular transport

We next investigated whether AT-LNTs function as conduits for direct material transport, a defining functional property of TNTs. To determine whether AT-LNTs provide a diffusion-accessible pathway for soluble molecules, we removed all membrane and actin dyes from the system, encapsulated the soluble fluorophore Alexa 488 inside actin-vesicles, and induced AT-LNT formation under standard conditions **(Fig. 5A)**. Alexa 488, initially confined within the vesicle lumen, readily diffused into the connected AT-LNTs and was detected throughout nearly all nanotube structures **(Fig. 5, B and C)**. These observations demonstrate that the lumen of AT-LNTs is continuously connected to the vesicle interior, forming hollow and enclosed membrane conduits that permit diffusion of soluble molecules along the nanotube axis.

**Figure 5.**
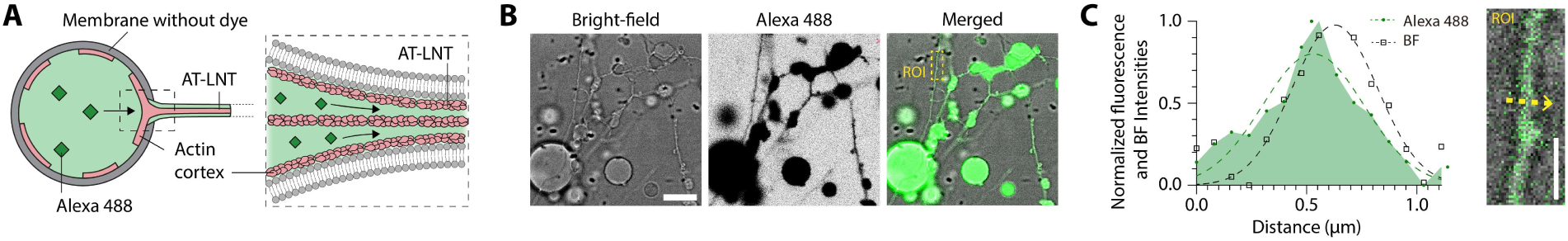
Solute diffusion along AT-LNTs. Actin-vesicles encapsulating Alexa 488, but without lipid and actin dyes, were triggered to form AT-LNTs in 20 mM Mg^2+^ buffer and observed under bright field and confocal fluorescence microscopy. **(A)** Schematics of an actin-vesicle encapsulating Alexa 488 and its diffusion into an AT-LNT. **(B)** Bright-field and confocal fluorescence microscopy images of Mg^2+^-triggered AT-LNTs from actin-vesicles encapsulating Alexa 488 (shown in inverted gray in the single-channel image and green in the merged image). Scale bar, 10 μm. **(C)** Normalized fluorescence and bright-field intensity profiles measured along a line perpendicular to the AT-LNT (yellow dashed arrow in the ROI inset). Scale bar, 2 μm.

We next examined whether AT-LNTs support intervesicular transport of the solutes. Co-culture of actin-vesicles with and without encapsulated Alexa 488 generated intervesicular AT-LNT connections **(fig. S15A)**. While lipid dyes readily redistributed across connected vesicles, transfer of soluble fluorophore into neighboring vesicles was observed only rarely and did not reach statistical significance **(fig. S15, B and C)**. These results indicate that lumenal continuity within AT-LNTs and the source vesicle alone is insufficient to establish efficient intervesicular solute exchange with the acceptor vesicle.

Efficient solute transfer would require complete membrane fusion between the AT-LNT terminus and the docked vesicle membrane, thereby establishing open lumenal continuity across both compartments. However, the membrane deformation energy required for such fusion is likely prohibitively high under the present experimental conditions (*44*, *45*), rendering stable intervesicular transport inefficient. Thus, while AT-LNTs reproduce the structural continuity and diffusion-accessible lumen characteristic of TNT-like conduits, they do not yet fully recapitulate the open-ended transport behavior observed in some cellular TNT systems (*7*, *9*, *46*).

### Bridging to living cells

We next investigated whether AT-LNTs formed from synthetic vesicle systems can extend beyond vesicle-vesicle connections to establish direct interfaces with living cells. Actin-vesicles were co-incubated with HEK293 cells, a commonly used model system for tunneling nanotube studies (*47*), and exposed to 20 mM Mg²⁺ buffer **(Fig. 6A)**. Under these conditions, AT-LNTs extended from actin-vesicles toward neighboring cells and established stable vesicle-cell connections.

**Figure 6.**
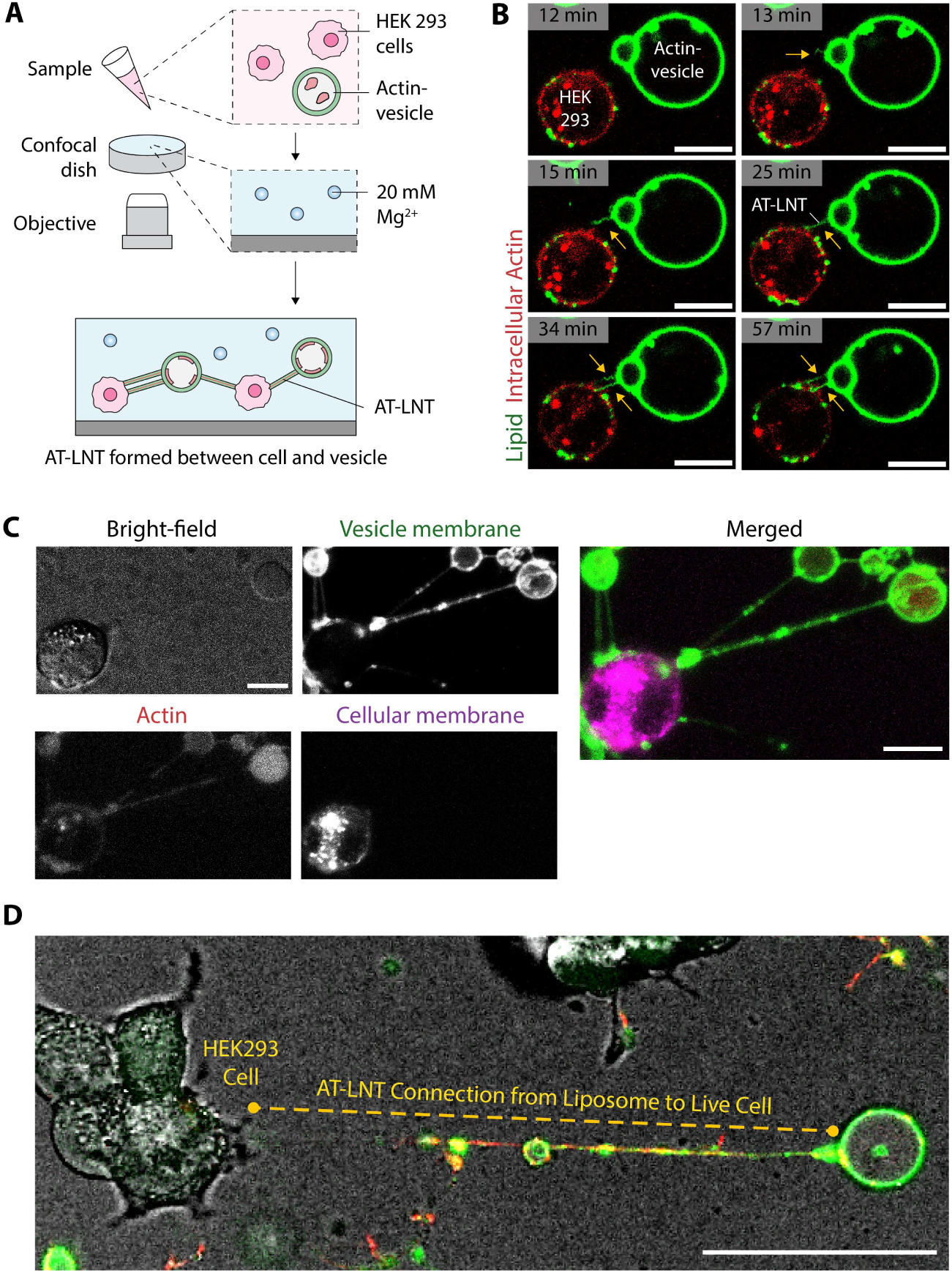
AT-LNT connection between actin-vesicles and live cell. **(A)** Schematic of the experimental design. Homogenously mixed actin-vesicles and HEK293 cells were introduced into 20 mM Mg²⁺ buffer. AT-LNTs developed from actin-vesicles and formed connections with HEK293 cells. **(B)** Time-lapse confocal fluorescence images showing AT-LNTs development (yellow arrows) between a HEK293 cell and an actin-vesicle. A neighboring but separated cell and actin-vesicle were observed (12 min), followed by initial AT-LNT development (13 min), elongation (15 min), docking onto a HEK293 cell (25 min), additional AT-LNT development (34 min), and further docking (57 min). Time indicates duration after Mg²⁺ exposure. Red, intracellular actin; green, vesicle membrane. Scale bars, 10 μm. See also movie S17. **(C)** AT-LNT connections between actin-vesicles and HEK293 cells observed by confocal fluorescence and bright-field microscopy. The merged image shows the vesicle membrane (green), intracellular and vesicle-encapsulated actin (red), and the cell membrane (magenta). Scale bars, 10 μm. **(D)** Merged confocal fluorescence and bright-filed image of a long AT-LNT connection (yellow dashed line) between actin-vesicle and HEK293 cells. Green, vesicle membrane; red, intracellular and vesicle-encapsulated actin. Scale bars, 50 μm.

Upon contact, cells exhibited active responses, including pulling on AT-LNTs and localized recruitment of endogenous actin to the docking sites **(Fig. 6B and movie S17)**. These responses indicate mechanical coupling between the synthetic nanotubes and the cellular cytoskeleton, suggesting that AT-LNTs can physically interface with active cellular membrane systems. The resulting vesicle-cell connections frequently spanned distances corresponding to several cell diameters and remained structurally stable over extended periods **(Fig. 6, C and D; movie S18)**.

We further assessed cell viability under the experimental conditions. Although the high Mg²⁺ and Na⁺-free environment is nonphysiological, most cells remained viable for several hours, sufficient for vesicle-cell connections to form **(fig. S16)**; AT-LNT connections persisted even during membrane blebbing and progressive cellular degradation, indicating substantial structural stability of the nanotube bridges.

The ability of AT-LNTs to directly bridge synthetic vesicles and living cells demonstrates that nanotubes generated in a minimal physicochemical system can establish stable interfaces with biological membranes. This finding extends the relevance of AT-LNTs beyond artificial vesicle networks and suggests that TNT-like nanotubes generated through minimal actin-membrane interactions may provide a versatile platform for probing intercellular connectivity across synthetic and living systems.

## Discussion

Cellular membrane tubules emerge through tightly coordinated interactions between lipid membranes and the actin cytoskeleton. Here, we demonstrate that TNT-like intervesicular nanotube connections can arise from a remarkably minimal physicochemical system composed only of lipid membranes, actin, and Mg²⁺. Upon Mg²⁺ exposure, actin-vesicles spontaneously generated actin bundle-embedded lipid nanotubes (AT-LNTs) that formed stable intervesicular connections resembling tunneling nanotubes (TNTs) **(Fig. 1)**. Systematic perturbation of membrane rigidity, membrane tension, Mg²⁺ concentration, ionic strength, and actin concentration revealed that AT-LNTs form only within a narrow physicochemical regime where membrane deformation and actin-membrane coupling are simultaneously permissive **(Fig. 2 and 3)**.

Unlike previous reconstitution systems that relied on membrane-bound actin nucleators (*18*, *19*), our system exploits direct electrostatic coupling between F-actin and PC membranes rendered positively charged through Mg²⁺ adsorption (*23–25*). Conditions that weakened actin-membrane association, including electrostatic repulsion by negatively charged membranes, low Mg²⁺ concentration, elevated ionic strength (*39*), or insufficient actin concentration, abolished AT-LNT formation **(Fig. 3)**, whereas excessive actin accumulation generated rigid actin cortices that suppressed membrane tubulation (*40*, *41*). These observations indicate that TNT-like nanotubes emerge only within an intermediate interaction regime in which actin-membrane coupling is sufficiently strong to support membrane protrusion while remaining permissive for membrane deformation.

AT-LNTs formed exclusively under conditions of low membrane deformation energy, namely liquid-disordered membrane phases and reduced membrane tension **(Fig. 2)**. A crude estimate of the equilibrium force required to extract a membrane tube with the average AT-LNT radius (∼30 nm; **Fig. 1G**) yields ∼17 pN for DOPC membranes but ∼86 pN for *Lo* phase membranes **(Supplementary Note 2)**, consistent with the selective AT-LNT formation from deformable *Ld* membranes. Considering the estimated 3-36 actin filaments within the nanotube lumen **(Supplementary Note 1)** and the reported actin polymerization force of ∼1 pN per filament (*48*, *49*), initial membrane protrusion by confined actin polymerization is mechanically plausible. However, classical actin-driven protrusion models alone cannot fully explain the longest AT-LNTs observed here, because membrane tension increases with tube elongation (*50*) and conventional mechanics predicts buckling of actin bundles at substantially shorter lengths (*11*). These observations suggest that long-range AT-LNT extension involves additional stabilization mechanisms beyond simple protrusive actin polymerization.

Our results further suggest that Mg²⁺-induced lipid nanotubes lacking internal actin (Mg-LNTs) represent precursor structures for AT-LNT formation. Mg-LNTs formed spontaneously from actin-free vesicles immediately after Mg²⁺ exposure **(movie S10)** and predominated under conditions that suppress actin-membrane association **(Fig. 3)**, indicating that membrane tubulation can initially be induced by Mg^2+^ independently of actin bundle formation. Together with the observed vesicle aggregation and hemifusion **(fig. S10A)**, these findings support a mechanism in which Mg²⁺ initiates membrane nanotubes, after which membrane-associated actin bundles stabilize and elongate the nanotubes that dock onto adjacent vesicles via Mg^2+^-mediated intermembrane adhesion (*36*-*38*) to form persistent intervesicular connections. This mechanism differs fundamentally from conventional actin-driven protrusion models in that membrane tubules appear to emerge prior to substantial actin bundle assembly within the lumen.

Importantly, AT-LNTs reproduced several defining structural and dynamic features of cellular TNTs. Cryo-EM analysis revealed lumenal actin bundles within the nanotubes **(Fig. 1F)**, while fluorescence imaging demonstrated long suspended intervesicular connections **(Fig. 1, B and C)**. Many AT-LNTs consisted of bundled individual nanotubes (iAT-LNTs) **(Fig. 4, B-D)**, analogous to bundled individual TNTs (iTNTs) observed in cells (*9*), and localized photodamage triggered progressive unwinding dynamics resembling the transition from bundled TNTs to single TNTs **(Fig. 4, E and F)** (*43*). The close correspondence between these behaviors suggests that higher-order TNT organization and remodeling can emerge from generic physical interactions between membrane nanotubes rather than requiring complex cellular regulatory machinery.

While soluble fluorophores readily diffused through the continuous lumen of AT-LNTs **(Fig. 5)**, efficient intervesicular transfer remained limited, likely because membrane fusion at nanotube-vesicle docking sites is energetically unfavorable. Nevertheless, AT-LNTs established direct interfaces between synthetic vesicles and living cells **(Fig. 6)**, indicating that nanotubes generated through minimal physicochemical interactions can interact with biological membranes. Importantly, these synthetic nanotubes spontaneously formed long-range vesicle-cell connections that closely resemble TNT-like cellular interactions, despite the complete absence of cellular regulatory machinery within the artificial vesicles. The observed cellular responses, including pulling forces and localized actin recruitment at docking sites **(Fig. 6B and movie S17)**, further suggest that AT-LNTs can mechanically couple with living cellular systems.

Although the present system does not yet demonstrate active signal or cargo transfer between artificial vesicles and cells, the spontaneous formation of TNT-like vesicle-cell bridges provides an important proof-of-concept that minimal synthetic membrane systems can directly interface with living cells through self-organized nanotubular structures. These findings therefore extend the significance of AT-LNTs beyond artificial membrane networks and suggest that minimal physicochemical interactions may capture fundamental aspects of TNT formation in living cells. More broadly, this system provides a promising framework for future studies aimed at engineering direct communication pathways between artificial and living cells, including signal exchange and targeted transport.

## Acknowledgments

Claude (Anthropic, Sonnet 5) was partially used for translation. The tool was not used to generate scientific content, analyses, interpretations, or conclusions. All scientific content was developed and verified by the authors, who take full responsibility for the final manuscript.

## Funding

National Research Foundation of Korea 2018R1A6A1A03024940 (SHP, SHL, CHK, KS)

National Research Foundation of Korea 2020R1A6C101A192 (SHP, SHL, CHK, KS)

NRF-A3 Foresight Program (KS)

Samsung Science and Technology Foundation SSTF-BA180112010 (KS)

## Author contributions

Conceptualization: SHP, LT, KS

Methodology: SHP, LT

Investigation: SHP, SHL, CHK, CY, XK

Visualization: SHP, SHL, CHK, CY, XK

Funding acquisition: KS

Project administration: KS

Supervision: LT, KS

Writing - original draft: SHP, SHL, KS

Writing - review & editing: SHP, LT, KS

## Data, code, and materials availability

All data are available in the main text or the supplementary materials.

## Supplementary Materials

## Materials and Methods

### Proteins and Reagents

Alpha-skeletal actin purified (> 99%) from rabbit skeletal muscle, fluorescent actin stain 670 phalloidin (Phal 670), and SiR-Actin kit containing verapamil and jasplakinolide conjugated to silicone rhodamine (SiR-actin) were purchased from Cytoskeleton, Inc. (Denver, CO). 1,2-dioleoyl-sn-glycero-3-phosphocholine (DOPC), 1,2-distearoyl-sn-glycero-3-phosphocholine (DSPC), cholesterol (ovine), 1,2-dioleoyl-sn-glycero-3-phospho-(1’-rac-glycerol) (DOPG), 1-palmitoyl-2-(dipyrrometheneboron difluoride)undecanoyl-sn-glycero-3-phosphocholine (TopFluor^TM^ PC), 23-(dipyrrometheneboron difluoride)-24-norcholesterol (TopFluor^TM^ Cholesterol), and 1,2-dioleoyl-sn-glycero-3-phosphoethanolamine-N-(lissamine rhodamine B sulfonyl) (18:1 Liss Rhod PE) were purchased from Avanti Polar Lipids, Inc. (Alabaster, AL). Mag-Fluo-4 (tetrapotassium salt), Alexa Fluor^TM^ 488 carboxylic acid (Alexa 488), Dulbecco’s Modified Eagle Medium (DMEM), fetal bovine serum (FBS), and penicillin-streptomycin-glutamine (100X), Vybrant^TM^ DiI cell-labeling solution, and LIVE/DEAD^TM^ cell imaging kit were purchased from Thermo Fisher Scientific (Waltham, MA). 2X DMEM was purchased from Welgene, Inc. (Gyeongsan, South Korea). Tris, sugar, glucose, adenosine 5′-triphosphate disodium salt (Na_2_ATP), MgCl_2_, NaCl, ethylenediaminetetraacetic acid (EDTA), chloroform, and magnesium ionophore I (Selectophore^TM^) were purchased from Sigma-Aldrich (Taufkirchen, Germany).

### Buffer Solutions

The standard internal buffer for actin-encapsulating vesicles consisted of 2 mM Tris-HCl (pH 7.8), 1 mM Na_2_ATP, 1 μM G-actin, 280 nM Phal 670, and sucrose added up to 300 mOsm/kg. Phal 670 was first added to tubes and dried under vacuum to remove methanol solvent before the addition of other solution components. The internal buffers were incubated for 12 h at 4 ℃ to ensure actin in a soluble monomeric state. For Mg^2+^ permeation assays, 15 μM Mag-Fluo-4 was additionally included. For experiments on solute diffusion along actin bundle-embedded lipid nanotubes (AT-LNTs,) 10 μM Alexa 488 was added instead. Actin concentrations were varied in control experiments to modulate the extent of actin–membrane interactions.

The standard external buffer for Mg^2+^-induced AT-LNT formation contained 10 mM Tris-HCl (pH 7.8), 20 mM MgCl_2_, and glucose adjusted up to 300 mOsm/kg. Glucose concentrations were varied to modulate osmolarity, MgCl_2_ concentrations were varied to examine Mg^2+^ effects, 2 mM EDTA was additionally included in the 0 mM Mg^2+^ condition, and NaCl was added to examine ionic strength effects.

All buffers were adjusted to pH 7.8 with 1 M HCl. Sucrose and glucose were added last, after all other components, to adjust osmolarity. Osmolarity was measured using a Fiske^®^ 210 Micro-Sample Osmometer (Advanced Instruments, Norwood, MA).

### Preparation of Actin-Encapsulating Vesicles (Actin-Vesicles)

All lipid stocks were dissolved in chloroform. The standard lipid mixture for actin-vesicles that generate AT-LNTs consisted of 94.5 mol% DOPC, 5 mol% Mg^2+^ ionophore I, and 0.5 mol% TopFluor^TM^ PC at a total lipid concentration of 2 mg/mL. For dual-color imaging of mixed actin-vesicles with differently labeled membranes, 0.5 mol% TopFluor^TM^ PC was replaced with 18:1 Liss Rhod PE. The lipid mixture for liquid-ordered (L_o_) phase PC vesicles consisted of 74.5 mol% DSPC, 20 mol% cholesterol, 5 mol% Mg^2+^ ionophore I, and 0.5 mol% TopFluor^TM^ cholesterol. The lipid mixture for liquid-disordered (L_d_) phase PG vesicles consisted of 94.5 mol% DOPG, 5 mol% Mg^2+^ ionophore I, and 0.5 mol% TopFluor^TM^ cholesterol. The lipid mixture for L_o_ PC and L_d_ PC phase-separated vesicles consisted of 38 mol% DOPC, 38 mol% DSPC, 18 mol% cholesterol, 5 mol% Mg^2+^ ionophore I, 0.5 mol% 18:1 Liss Rhod PE, and 0.5 mol% TopFluor^TM^ cholesterol. The lipid mixture for L_o_ PC and L_d_ PG phase-separated vesicles consisted of 38 mol% DSPC, 38 mol% DOPG, 18 mol% cholesterol, 5 mol% Mg^2+^ ionophore I, 0.5 mol% 18:1 Liss Rhod PE, and 0.5 mol% TopFluor^TM^ cholesterol.

Actin-vesicles were formed by the electroformation method (*51*). A 40-μL aliquot of each lipid mixture was spread on the conductive side of two indium tin oxide (ITO)-coated glasses (16 cm^2^ area, 30 Ω; Fine Chemicals Industry, Seoul, South Korea) and dried under vacuum for 1 h to form lipid films. The ITO glasses with lipid films were attached to each side of a 1 mm-thick silicone rubber spacer with a 250 μL central chamber. Lipid films were hydrated with the prepared internal buffer, and a 10 Hz sinusoidal wave at 500 mV_RMS_ was applied for 2.5 h using a function generator (Tektronix Inc., Beaverton, OR, USA). Actin-vesicles were kept at 4 ℃ and used in experiments on the day of preparation.

### Mg^2+^-Triggered Formation of AT-LNTs from Actin-Vesicles

A confocal imaging dish with a polystyrene plastic window (PS/Flux; 13-mm diameter; SPL Life Sciences, Pocheon, South Korea) was used to prevent strong adhesion of vesicles to the substrate under high divalent cation concentrations. External Mg^2+^ buffer (180 μL) was first loaded into the confocal dish. Subsequently, a small aliquot (2.5 μL) of actin-vesicles was gently transferred into the external buffer **(fig. S1A)** to trigger AT-LNT formation.

### Fluorescence and Optical Microscopy

Bright-field (BF), differential interference contrast (DIC), fluorescence, and confocal fluorescence laser microscopy images were acquired using a Leica TCS SP8 confocal microscope (Leica-Microsystems, Mannheim, Germany). For confocal fluorescence imaging, TopFluor^TM^ PC, TopFluor^TM^ cholesterol, Alexa 488, and Mag-Fluo-4 were excited at 499 nm with 1% laser power, and 18:1 Liss Rhod PE and DiI lipophilic membrane stain were excited at 555 nm with 1% laser power. Phal 670 and SiR-actin were excited at 640 nm at 2.5% laser power. Images were acquired using 40× water and 60 × oil immersion lens with the pinholes set at 1 or 2 AU.

### Cryo-Electron Microscopy

A 2.5-μL aliquot of external buffer was applied to glow-discharged, carbon-coated holey grids (Quantifoil, Großlöbichau, Germany). Subsequently, 0.5 μL of actin-vesicles was added to the applied external buffer on the grid and incubated for 15 min to allow AT-LNT formation. The sample-loaded grids were blotted from the back side with filter paper for 7 s and vitrified by rapid plunging into liquid ethane. Electron micrographs were acquired using a JEM-2100Plus (JEOL Ltd., Tokyo, Japan) or a Glacios 200 kV Cryo-TEM (Thermo Fisher Scientific, Waltham, MA).

### Cell Experiments

HEK293 cells were cultured in complete medium consisting of DMEM supplemented with 10% (v/v) FBS and 1% (v/v) penicillin-streptomycin-glutamine (100X) at 37 °C in a humidified incubator with 5% CO₂.

Intracellular actin was stained by incubating adherent cells with 500 nM SiR-actin and 5 nM verapamil in complete medium for 4 h at 37 °C and 5% CO₂. Cellular membranes were stained by incubating the cells with 5 μM DiI cell-labeling solution in complete medium for 20 min at 37 °C and 5% CO₂. The dye-containing medium was removed, and the cells were washed three times with complete medium. The cells were then maintained in suspension before experiments.

For the cell viability assay of the HEK293 cells in standard 20 mM Mg^2+^ external buffer, cells were suspended in a 1:1 mixture of 2X LIVE/DEAD^TM^ Cell Imaging Kit working solution and 2X external buffer (20 mM Tris-HCl, 40 mM Mg^2+^, glucose adjusted to 700 mOsm/kg). The cell suspension was transferred into a confocal dish and incubated at 37 °C with 5% CO₂ before observation by confocal microscopy.

For vesicle–cell AT-LNT connection formation, 2 μL of stained cell suspension (approximately 80,000 cells) was gently mixed with 20 μL of actin-vesicles. The mixture was then loaded into standard Mg^2+^ external buffer adjusted to an osmolarity of 350 mOsm/kg, close to that of complete medium **(Fig. 6A)**.

### Statistics

Data are presented as mean ± SD. Statistical significance between data sets was determined using Student’s *t* test, and *p* values were considered nonsignificant when *p* ≥ 0.05. Correlations between paired data sets were evaluated using Pearson’s correlation analysis and were considered correlated when the absolute value of Pearson correlation coefficient was |*r*| > 0.5.

## Supplementary Text

<u>Supplementary Note 1: Estimation of the number of constituent filaments and persistence length of an actin bundle inside an AT-LNT</u>

Given the known diameter of an actin filament (*D*_filament_ = 7 *nm*) and measured diameter of actin bundles (*D*_bundle_ = 13 − 42 *nm*) within AT-LNTs **(Fig. 1, F and G)**, the number of constituent actin filaments (*N*_filaments_) can be estimated as,

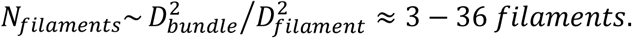

Assuming that Mg^2+^-crosslinked actin filaments can freely slide relative to one another (*52*), the persistence length of the internal actin bundles 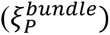 is given by,

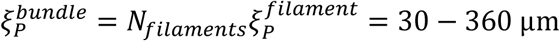

where the known persistence length of an actin filament 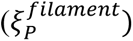 is approximately 10 μm (*53*, *54*)

<u>Supplementary Note 2: Estimation of Required Force for AT-LNT Generation</u>

The conventional equilibrium tether force for a membrane tube (*29*, *30*) is given by:

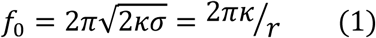

where n is the bending rigidity of the membrane, *σ* is the membrane tension, and *r* is the radius of an extracted tube.

Substituting the known values of bending rigidities for a DOPC vesicle (κ ≈ 20 k_B_T) (*33*) and L_o_ phase (κ ≈ 100 k_B_T) (*32*) vesicle into **Eq. 1**, the equilibrium forces required to extract the tube with a radius of 30 nm are approximately 17 pN and 86 pN, respectively.

On the other hand, when a membrane tube is extracted long enough from a vesicle adhered to a substrate and the membrane tension is not controlled by micropipette aspiration, a more precise relation (*50*) between the extracted length (*L*) of a tube and the required pulling force (*f*) is given by:

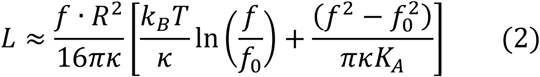

where *R* is the vesicle radius and *K*_A_ is stretch modulus of the membrane.

Substituting the values above (*f*_0_ ≈ 17 *pN*) and the known stretch modulus for a DOPC vesicle (*K*_A_ ≈ 250 mN m⁻¹) (*33*) into **Eq. 2**, the force required to extract a tube of radius 30 nm and length 20 µm from a vesicle of radius 5 µm is approximately 45 pN.

---

**Fig. S1.**
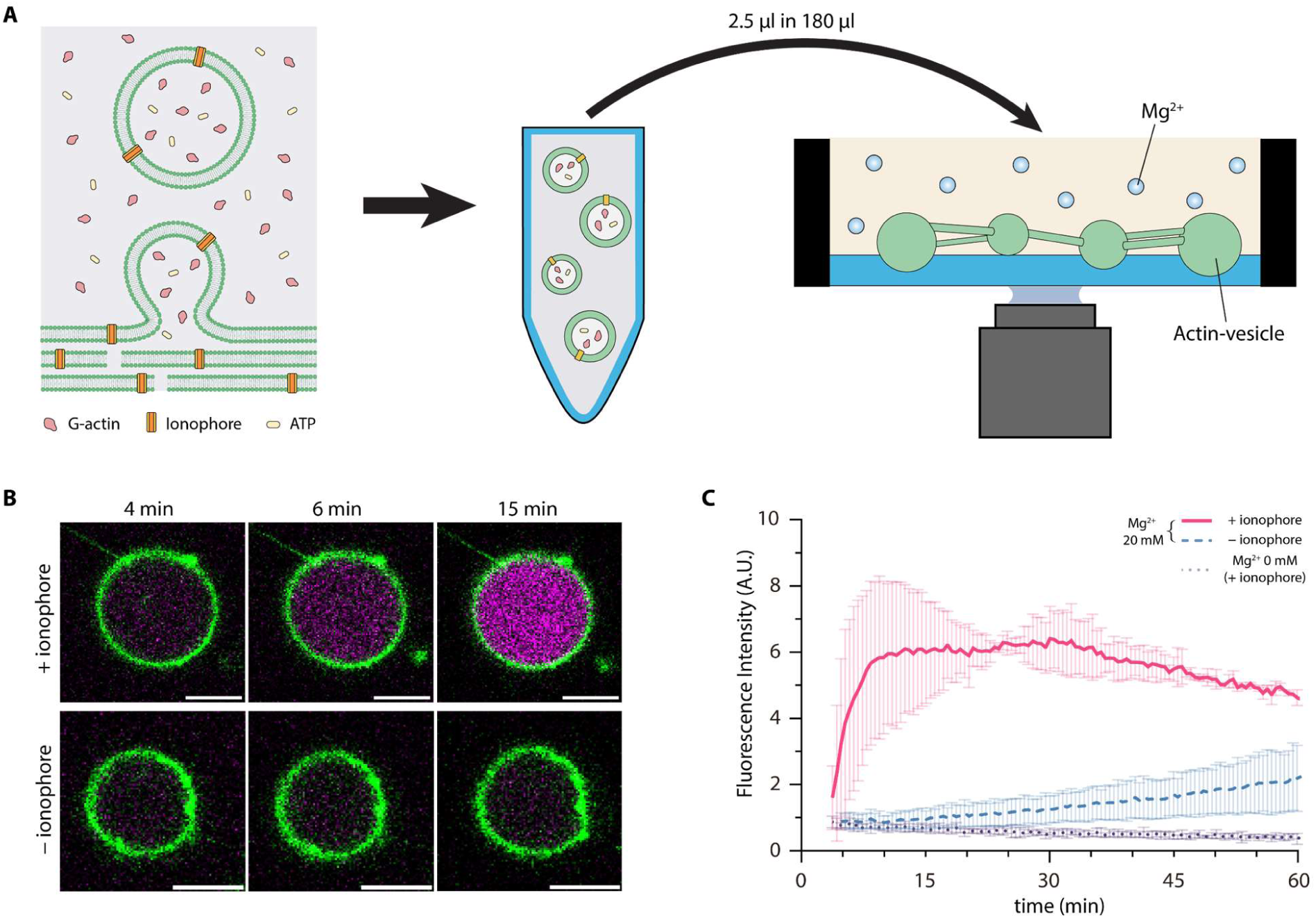
Mg^2+^ influx into actin-vesicles via Mg^2+^ ionophore. **(A)** Experimental workflow. Actin-vesicles (1 μM actin) with Mg^2+^ ionophore-embedded membrane were prepared by electroformation. Vesicles were added to 20 mM Mg^2+^ external buffer and observed under microscopes. **(B)** Representative time-lapse confocal fluorescence images showing Mg^2+^ influx. Actin-vesicles containing Mg^2+^ indicator (magenta) with lipid membrane labeled (green) were prepared with and without Mg^2+^ ionophore and introduced into external Mg^2+^ buffer at t = 0. Scale bars, 10 μm. **(C)** Quantification of Mg²⁺ indicator fluorescence intensity in actin-vesicles after exposure to external Mg²⁺ buffer. Data represent mean ± SD (n = 5 vesicles per condition).

**Fig. S2.**
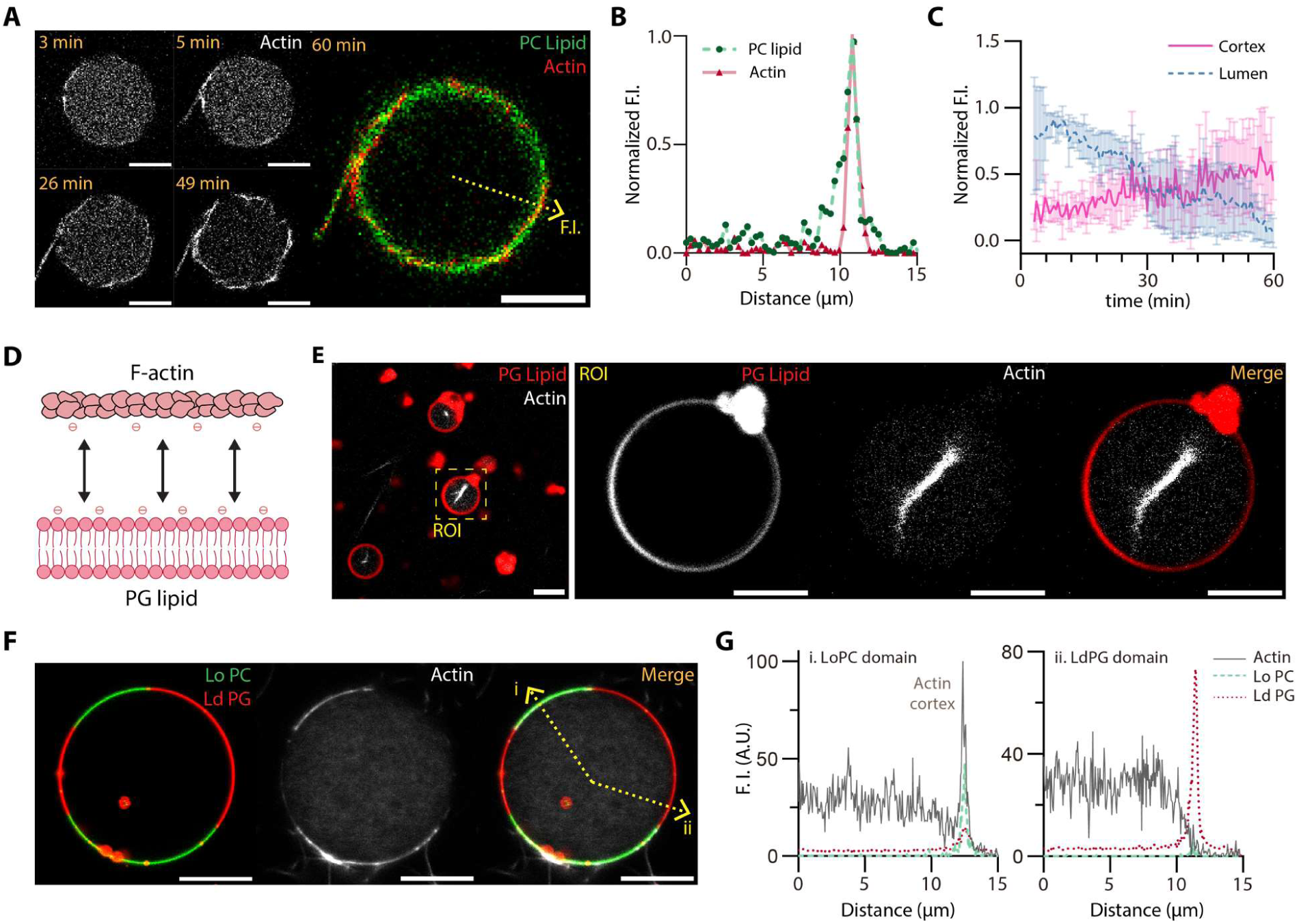
Actin–membrane interactions varied by membrane charge. **(A)** Representative time-lapse confocal fluorescence images of actin (gray) inside DOPC vesicles following transfer into external Mg^2+^ buffer at t = 0, imaged at 3, 5, 26, 49, 60 min. The merged image at 60 min shows actin (red) and lipid membrane (green). Scale bars, 10 μm. **(B)** Normalized fluorescence intensity profiles of lipid (green dashed) and actin (red) measured along the yellow dashed arrow in (A). **(C)** Mean lumenal (blue dashed) and maximal cortical (magenta) actin normalized fluorescence intensities in vesicles over time. Data represent mean ± SD (n = 5 vesicles). **(D)** Schematic illustrating electrostatic repulsion between negatively charged F-actin and the negatively charged PG membrane. **(E)** Representative confocal fluorescence images of actin-vesicles of DOPG membrane (red) under standard conditions, showing actin (gray) bundles localized at the lumenal center. Scale bars, 10 μm (left) and 5 μm (right). **(F)** Representative confocal fluorescence images of actin (gray) encapsulated in a Lo PC (green) and Ld PG (red) phase-separated vesicle. Scale bar, 10 μm. **(G)** Fluorescence intensity profiles of PC lipid (green dashed) and PG lipid (red dashed) and actin (gray) measured across the yellow dashed arrow in (F), for **(i)** the Lo PC domain and **(ii)** the Ld PG domain.

**Fig. S3.**
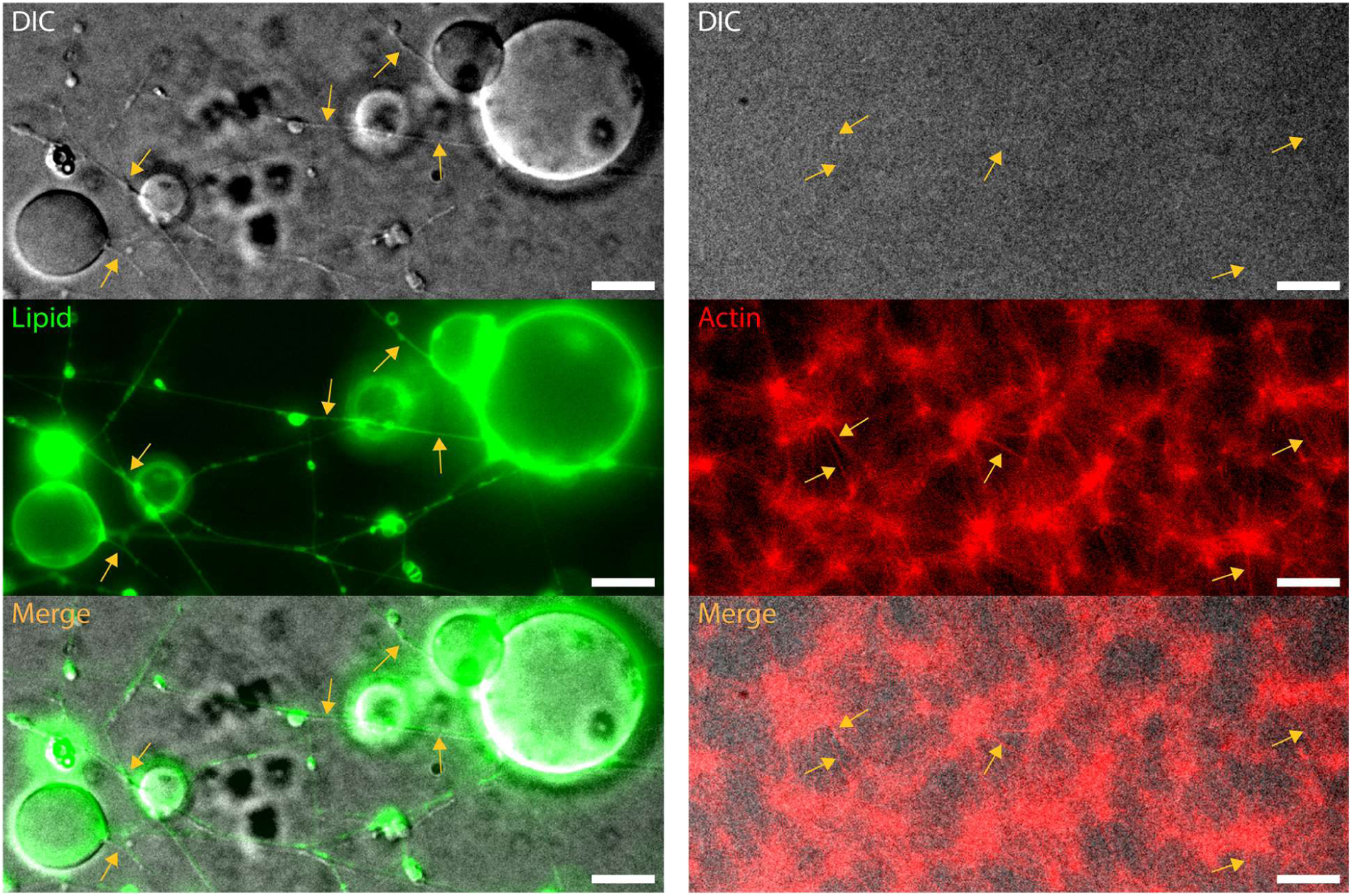
AT-LNTs and actin bundles distinguished by differential interference contrast (DIC) microscopy. Representative DIC and fluorescence images of membrane tubes (left) and actin bundles (right). **Left**: Membrane tubes formed from actin-vesicles under the standard conditions were imaged by DIC and fluorescence microscopy (green, lipid membrane). Membrane tubes are visible in DIC images (yellow arrows). **Right**: Actin bundles formed by incubating 10 μM actin in 20 mM Mg^2+^ buffer are not visible in DIC (yellow arrows indicate actin bundles identified by fluorescence). Scale bars, 10 μm.

**Fig. S4.**
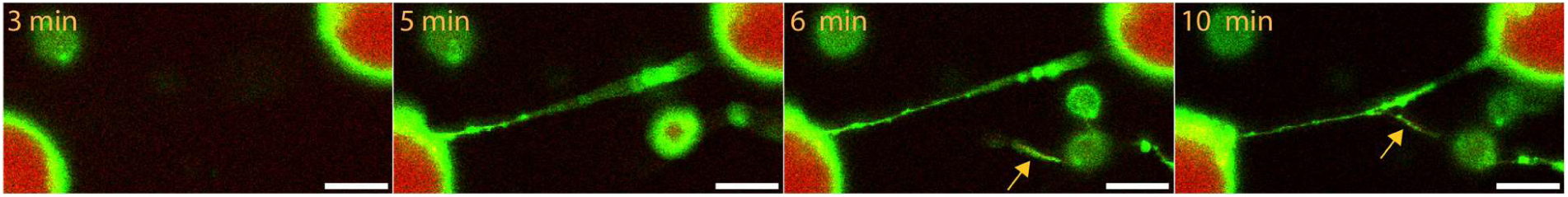
Formation of an AT-LNT branch through inosculation. Representative time-lapse confocal fluorescence images of AT-LNTs (green, lipid membrane; red, actin). AT-LNT formation was triggered by transferring into 20 mM Mg^2+^ buffer at t = 0. A branched AT-LNT connection formed as a growing AT-LNT (visible at 6 min, yellow arrow) inosculated with a preexisting AT-LNT (10 min, yellow arrow). Scale bar, 5 μm. See also movie S3.

**Fig. S5.**
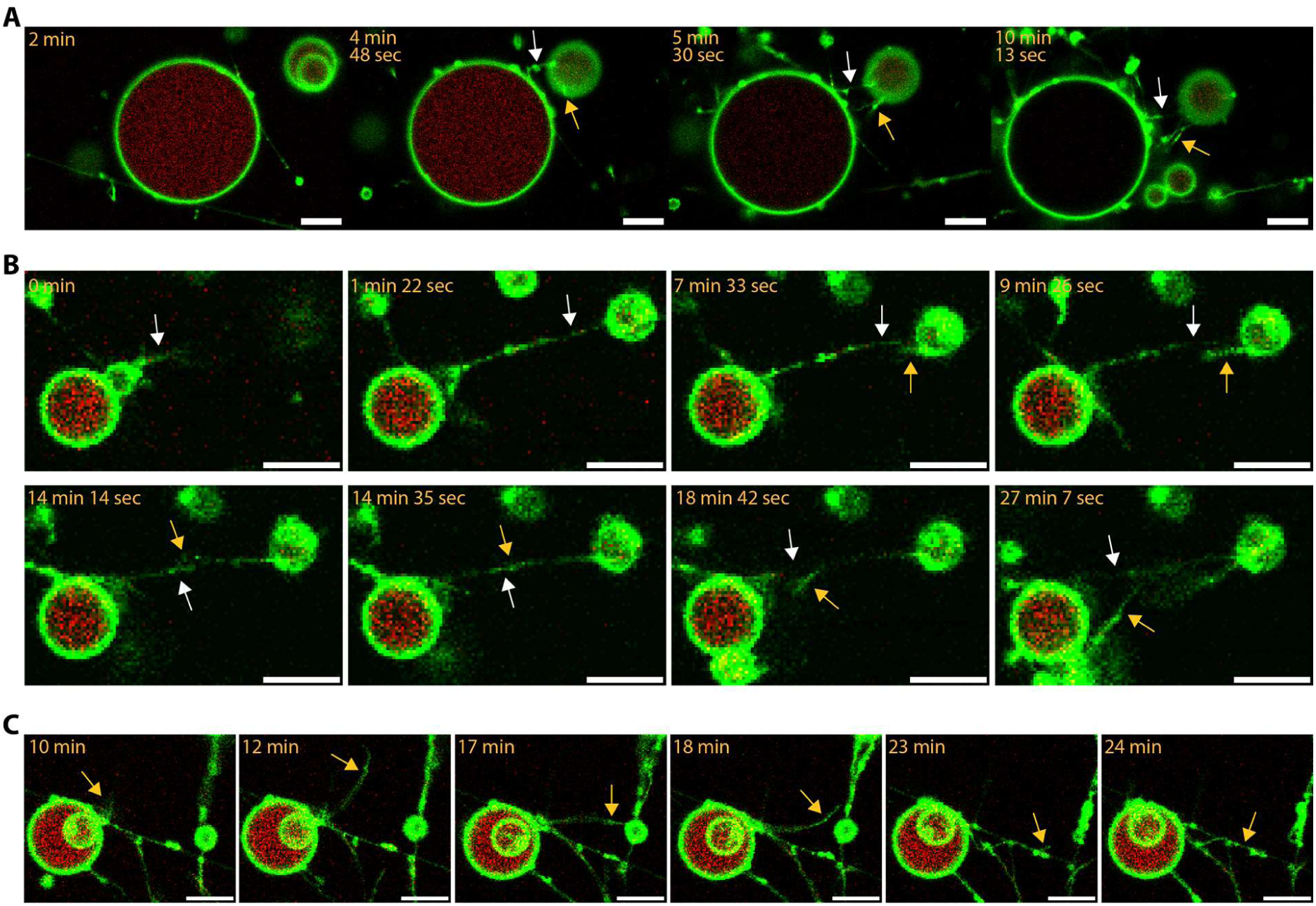
Additional growth of AT-LNTs at docking/growth sites developing into bundle and branches. Representative time-lapse confocal fluorescence images of AT-LNTs (green, lipid membrane; red, actin) **(A)** Multiple AT-LNTs forming at growth and docking sites. The first AT-LNT connection (white arrow) formed (at t = 4 min 48 sec), new AT-LNTs (yellow arrow) developed near initial growth site (t = 5 min 30 sec), and subsequently elongated to dock onto the connected vesicle (t = 10 min 13 sec). Time indicates duration from Mg^2+^ exposure. Scale bar, 10 μm. See also movie S4. **(B)** Bundling of AT-LNTs. An initial AT-LNT (white arrow) developed and docked onto a neighboring actin-vesicle (t = 1 min 22 sec). A new AT-LNT (yellow arrow) subsequently developed (t = 7 min 33 sec), elongated (t = 9 min 26 sec), bundled with the initial AT-LNT (t = 14 min 14 sec to 18 min 42 sec), and docked onto the counterpart actin-vesicle (t = 27 min 7 sec). Time indicates duration after imaging began. Scale bar, 10 μm. See also movie S5. **(C)** Branched configuration formed by an additional AT-LNT originating from a growth/docking site. An additional AT-LNT (yellow arrow) developed (t = 10 min), elongated (t = 12 min), docked onto a neighboring vesicle (t = 17 min), detached (t = 18 min), and inosculated with a preexisting AT-LNT (t = 23–24 min) to form a branched configuration. Time indicates duration after Mg^2+^ exposure. Scale bar, 10 μm. See also movie S6.

**Fig. S6.**
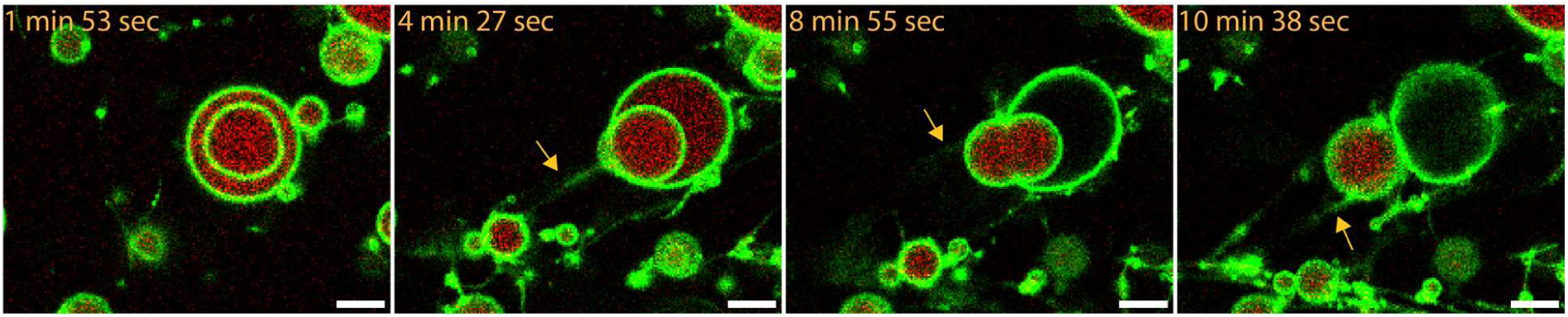
Formation of intervesicular AT-LNT connections via vesicle escape. A smaller vesicle was preformed inside a larger vesicle (t = 1 min 53 sec). The internal vesicle attached to the inner leaflet of the larger vesicle, developed an AT-LNT that connects to a preexisting AT-LNT outside (yellow arrow; t = 4 min 27 sec), and then subsequently escaped from the larger vesicle (t = 8 min 55 sec–10 min 38 sec). Time indicates duration after imaging began. Green, lipid; red, actin. Scale bar, 10 μm. See also movie S7.

**Fig. S7.**
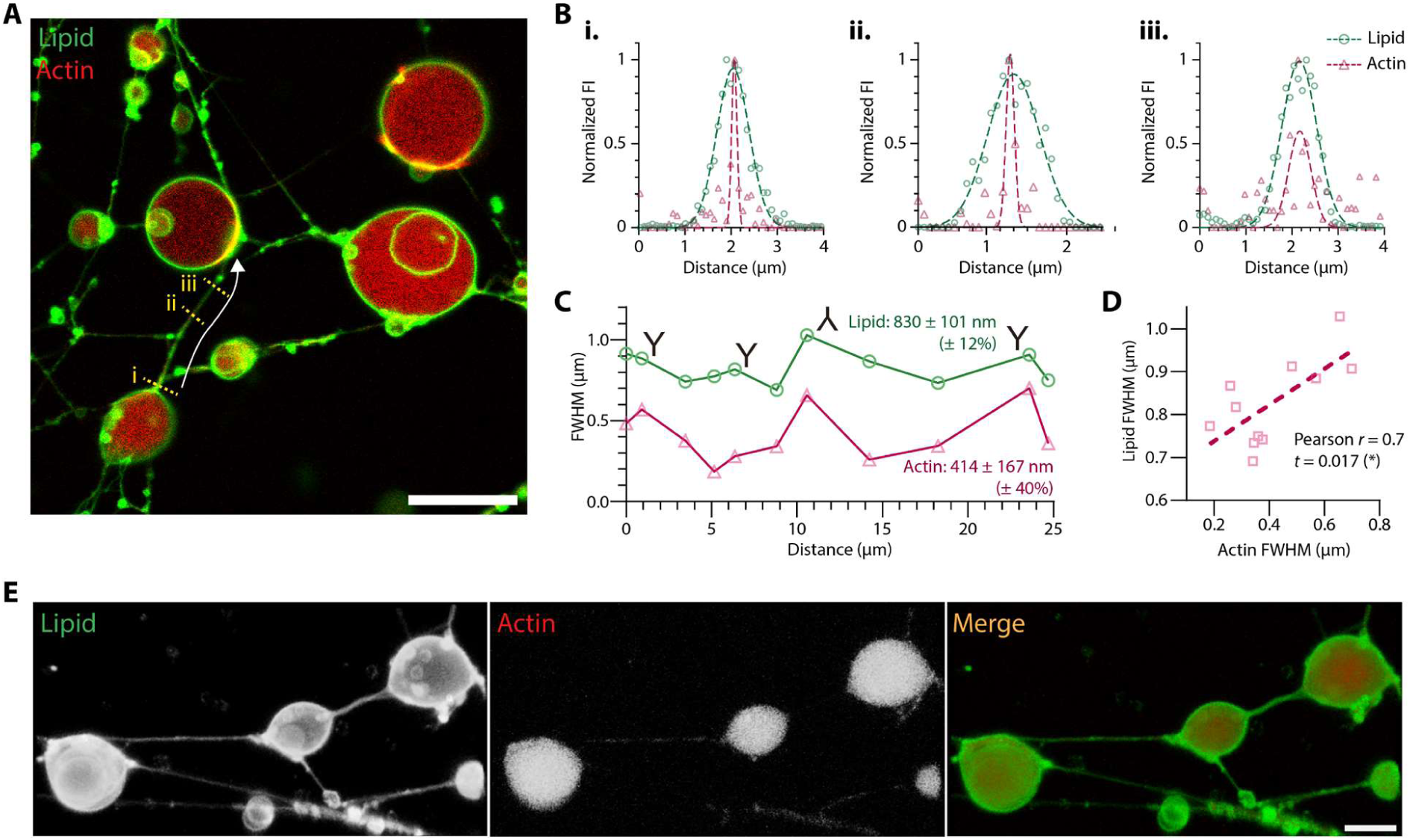
Actin distribution inside and along LNTs. Representative confocal fluorescence images of AT-LNTs under standard conditions (green, lipid membrane; red, actin) **(A)** Intervesicular AT-LNT connections, Scale bar, 10 μm. **(B)** Additional normalized fluorescence intensity profiles of lipid and actin measured along the yellow dashed lines **(i, ii, iii)** in (A). **(C)** Full width at half maximum (FWHM) of lipid and actin signals measured at 11 points along the intervesicular AT-LNT connection in the direction of the white arrow in (A). **“Y”** and **“inverted Y”** represent the positions of diverging and converging branches, respectively. **(D)** Pearson’s correlation coefficient between paired FWHM values of lipid and actin at each data point in (C). **(E)** 3D reconstruction of intervesicular AT-LNT connections generated from Z-stacks of confocal fluorescence microscopy images, viewed from above. Scale bar, 20 μm. See also movie S9 for the full rotation and sectional view of the 3D reconstruction.

**Fig. S8.**
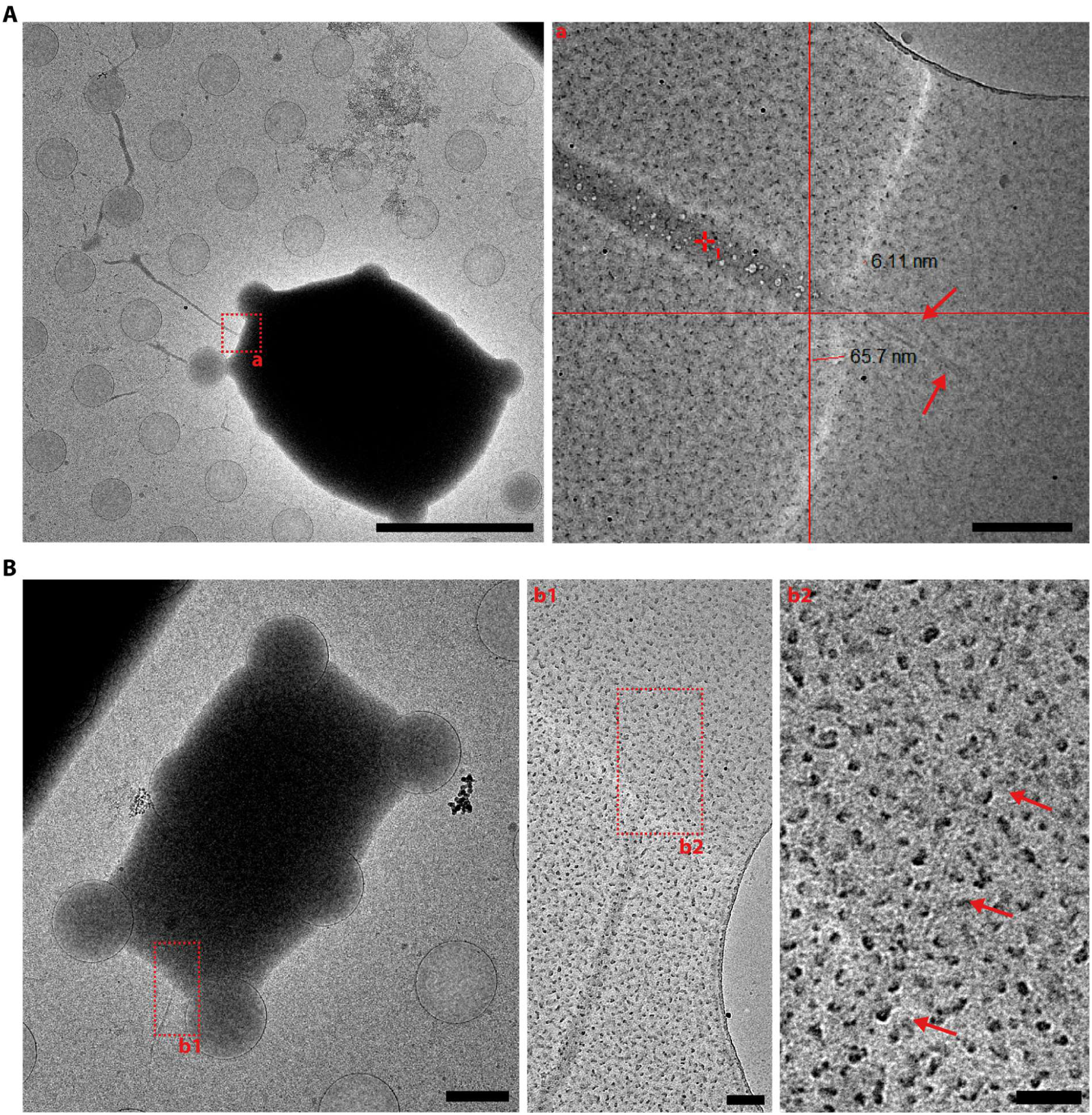
Cryo-EM images of AT-LNT growth or docking sites. Representative cryo-EM images of AT-LNTs induced under the standard conditions. **(A)** Fragmented actin filaments (red arrows) observed at a growth/docking site. Scale bars, 5 μm (left) and 200 nm (inset a, right). **(B)** An actin bundle (red arrows) extending from the lumenal interior of an actin-vesicle into an AT-LNT. Scale bars, 1 μm (left), 100 nm (inset b1, middle), and 50 nm (inset b2, right).

**Fig. S9.**
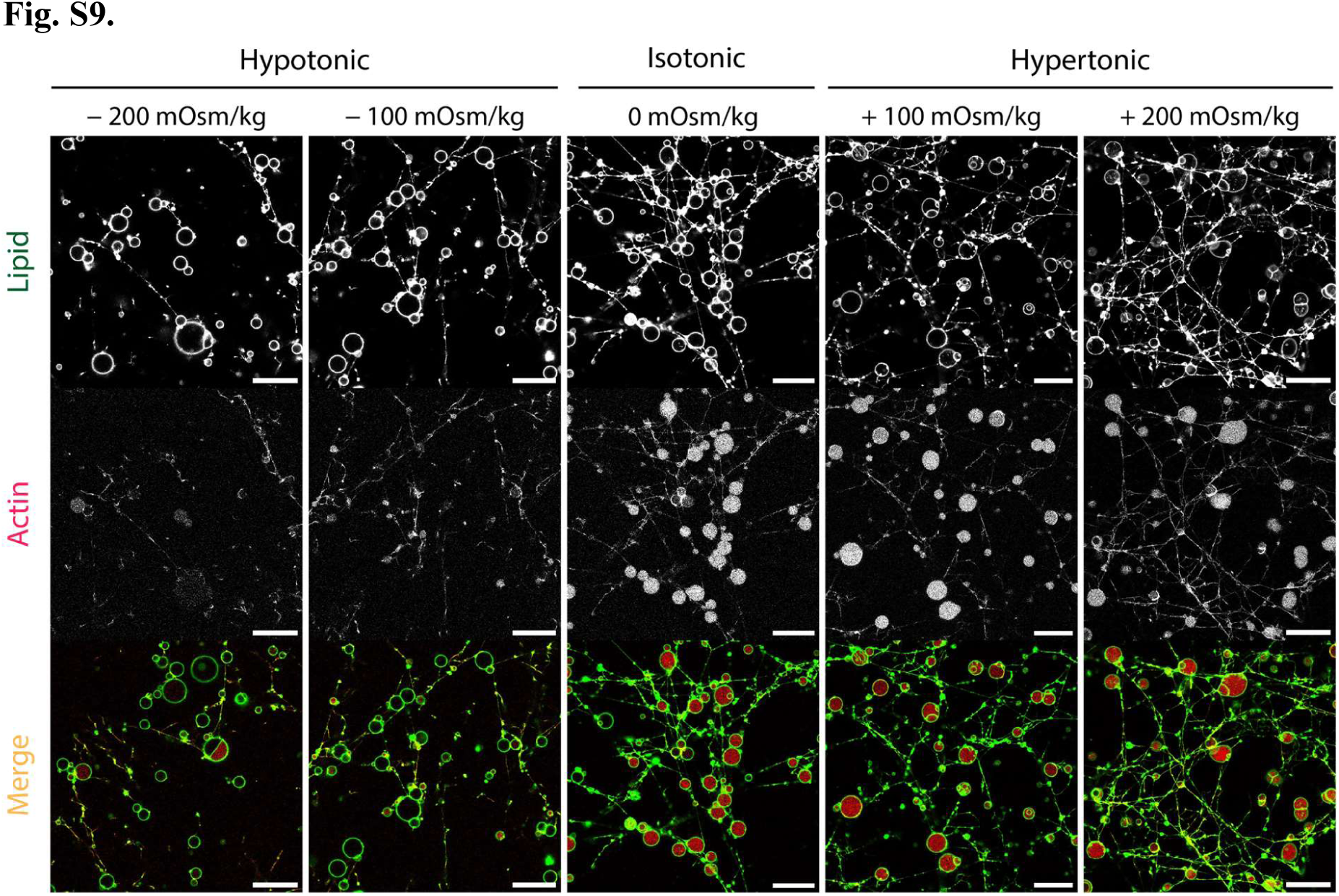
AT-LNT formation under different osmotic conditions. Representative confocal fluorescence images of AT-LNTs formed in 20 mM Mg²⁺ buffer at different osmotic conditions. Leakage of actin and fragmentation of lipid membranes were observed under hypoosmotic conditions (−100 and −200 mOsm/kg). More extensive intervesicular AT-LNT networks were observed under hyperosmotic conditions (+100 and +200 mOsm/kg). Green, lipid; red, actin. Scale bars, 50 μm.

**Fig. S10.**
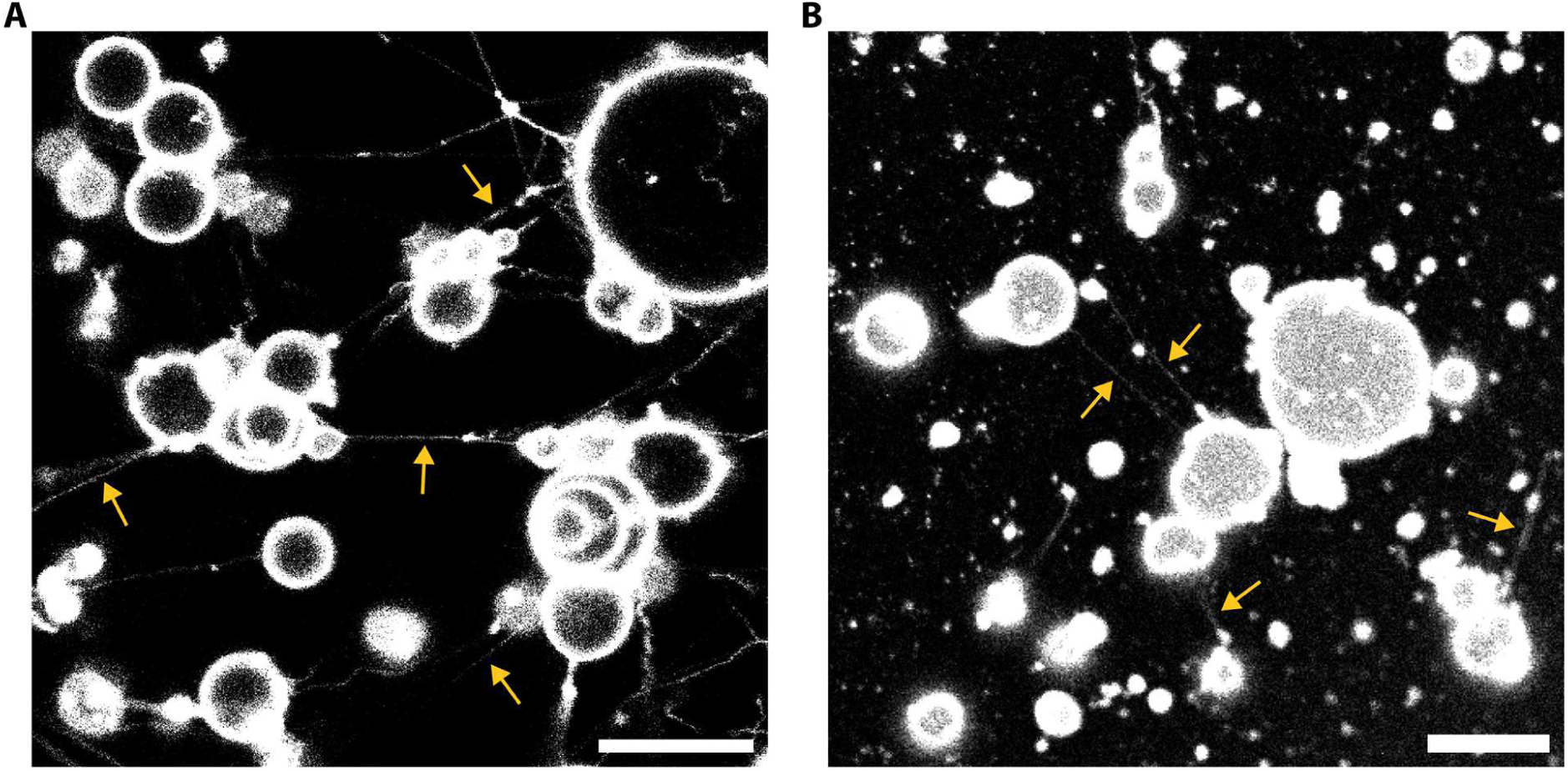
Mg²⁺-triggered LNT formation from DOPC vesicles without actin. Representative confocal fluorescence images of DOPC vesicles without actin in 20 mM Mg^2+^ buffer. Gray, lipid membrane. **(A)** Thermally fluctuating LNTs (yellow arrows) are spontaneously generated from DOPC vesicles and connect to neighboring vesicles. The majority of vesicles appear aggregated and hemifused with one another. Scale bar, 20 μm. **(B)** Vesicles and LNTs (yellow arrows) are observed adhered to the substrate. Scale bar, 20 μm. See also movie S10.

**Fig. S11.**
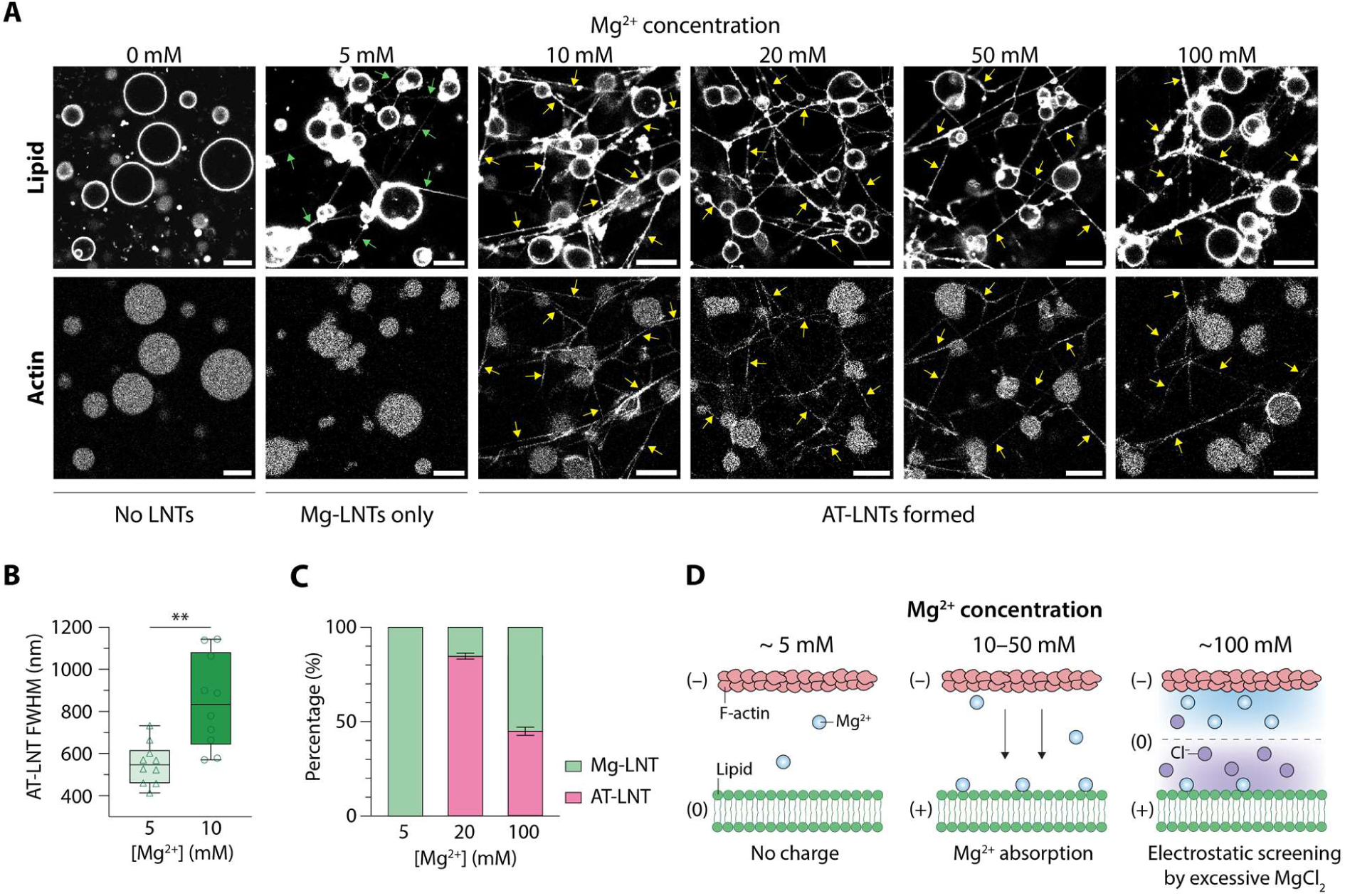
Mg^2+^ concentrations control AT-LNT formation. **(A)** Representative confocal fluorescence images of Mg^2+^-induced lipid nanotubes (Mg-LNTs; green arrows) and AT-LNTs (yellow arrows) formed from actin-vesicles (1 μM actin) at varying Mg^2+^ concentrations (in Na+-free external buffer). Scale bars, 10 μm. **(B)** Cross-sectional FWHMs of lipid signals comparing Mg-LNTs (5 mM Mg^2+^) and AT-LNTs (10 mM Mg^2+^) (n = 10 nanotubes per condition; data represent mean ± SD). **(C)** Proportion of Mg-LNTs and AT-LNTs at different Mg^2+^ concentrations. Data represent mean ± SD; n = 3 images (387.5 μm × 387.5 μm) per condition. **(D)** Schematics showing Mg^2+^-mediated actin–membrane interactions at varying Mg^2+^ concentrations.

**Fig. S12.**
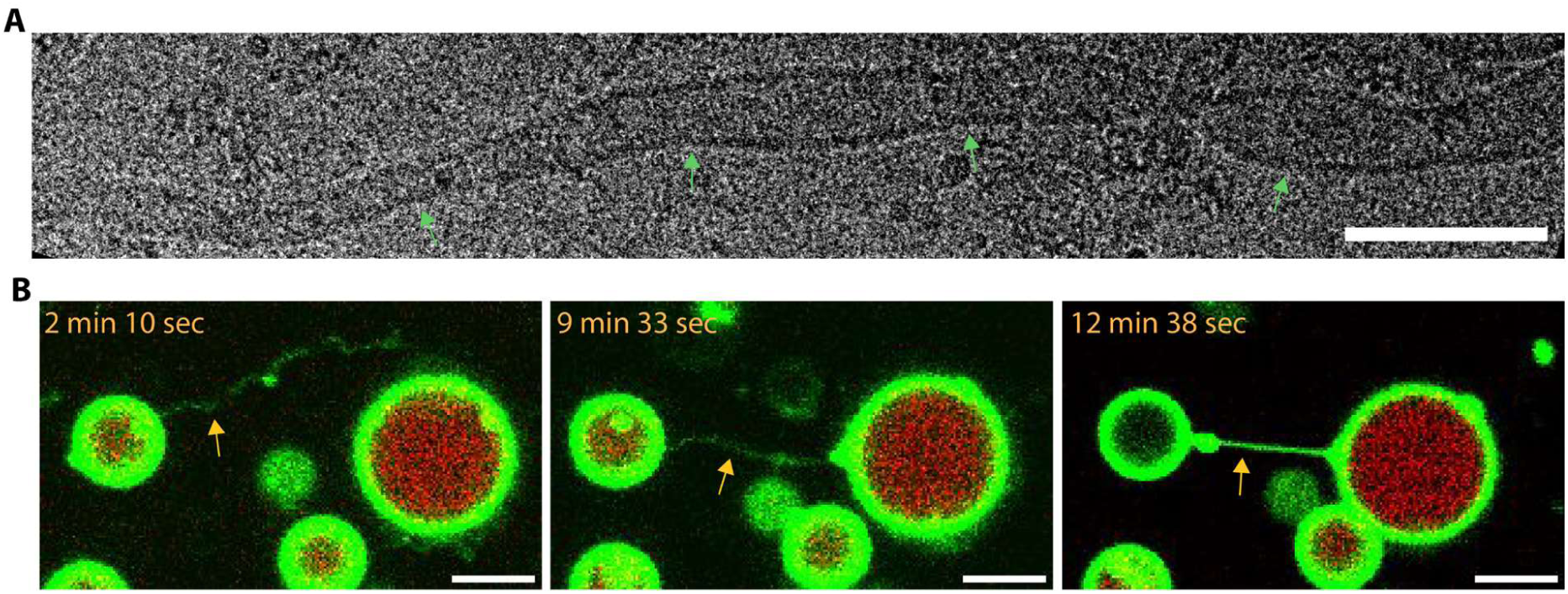
LNTs without internal actin formed under high ionic strength. **(A)** Cryo-EM image of an LNT formed at 150 mM ionic strength (90 mM Na⁺, 20 mM Mg²⁺, 1 μM actin). Green arrows indicate the lipid bilayer. Scale bar, 100 nm. **(B)** Time-lapse confocal fluorescence imaging of intervesicular Mg-LNT formation (yellow arrows) from actin-vesicles (1 μM actin) at 150 mM ionic strength (90 mM Na⁺, 20 mM Mg²⁺). Time indicates duration after imaging began. Green, lipid; red, actin. Scale bars, 10 μm. See also movie S14.

**Fig. S13.**
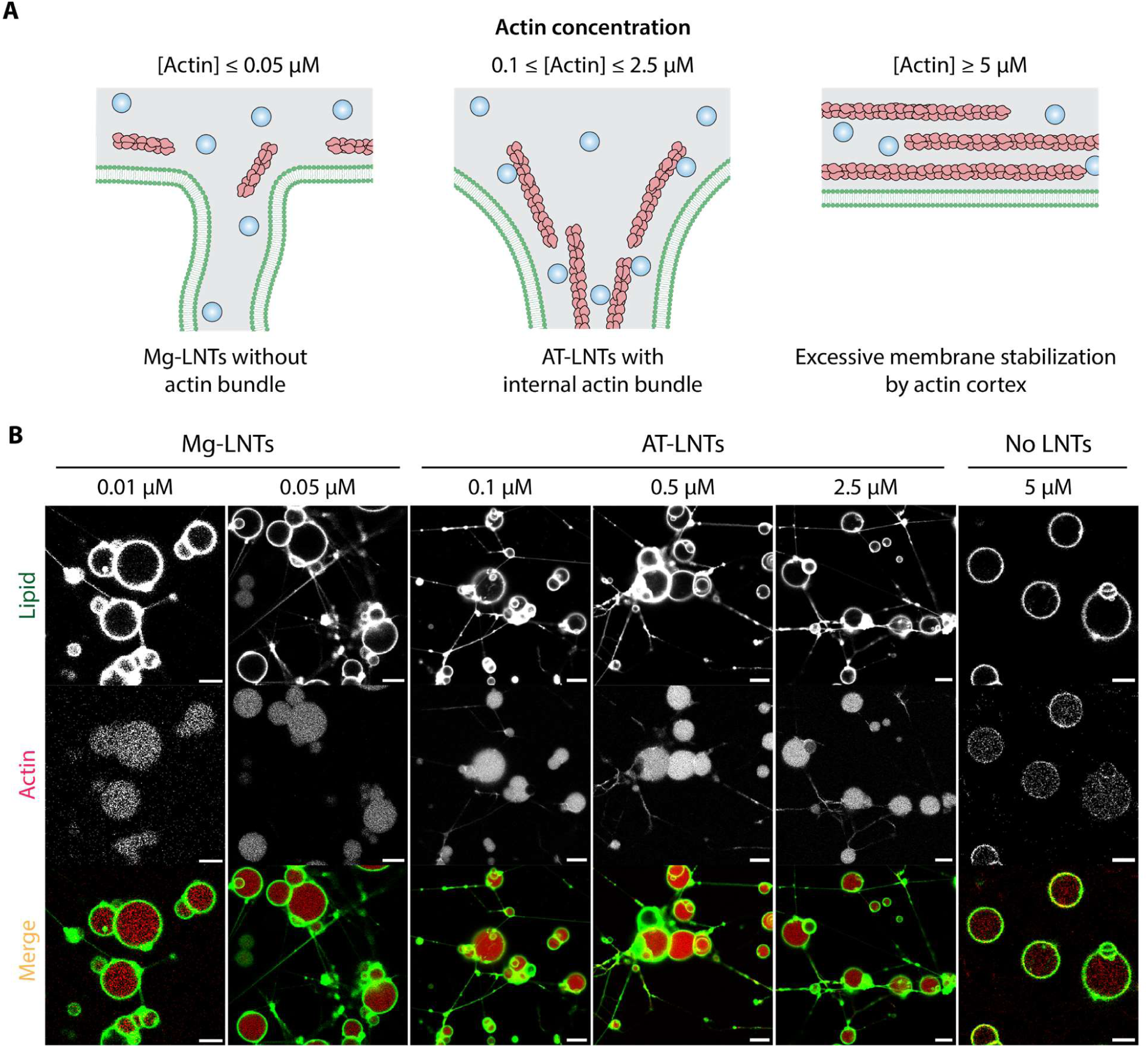
Actin concentration controls AT-LNT formation. **(A)** Schematics showing Mg-LNT formation at low actin concentrations (left), AT-LNT formation at optimal actin concentrations (center), and inhibition of LNT formation by excessive membrane stabilization at high actin concentrations (right). **(B)** Representative confocal fluorescence images of actin-vesicles with varying actin concentration in 20 mM Mg²⁺ buffer (Na^+^-free). Mg-LNTs without actin colocalization formed at actin concentrations of 0.01 and 0.05 μM. AT-LNTs with actin colocalization formed at 0.1, 0.5, and 2.5 μM actin. No LNTs formed at 5 μM actin. Green, lipid; red, actin. Scale bars, 10 μm.

**Fig. S14.**
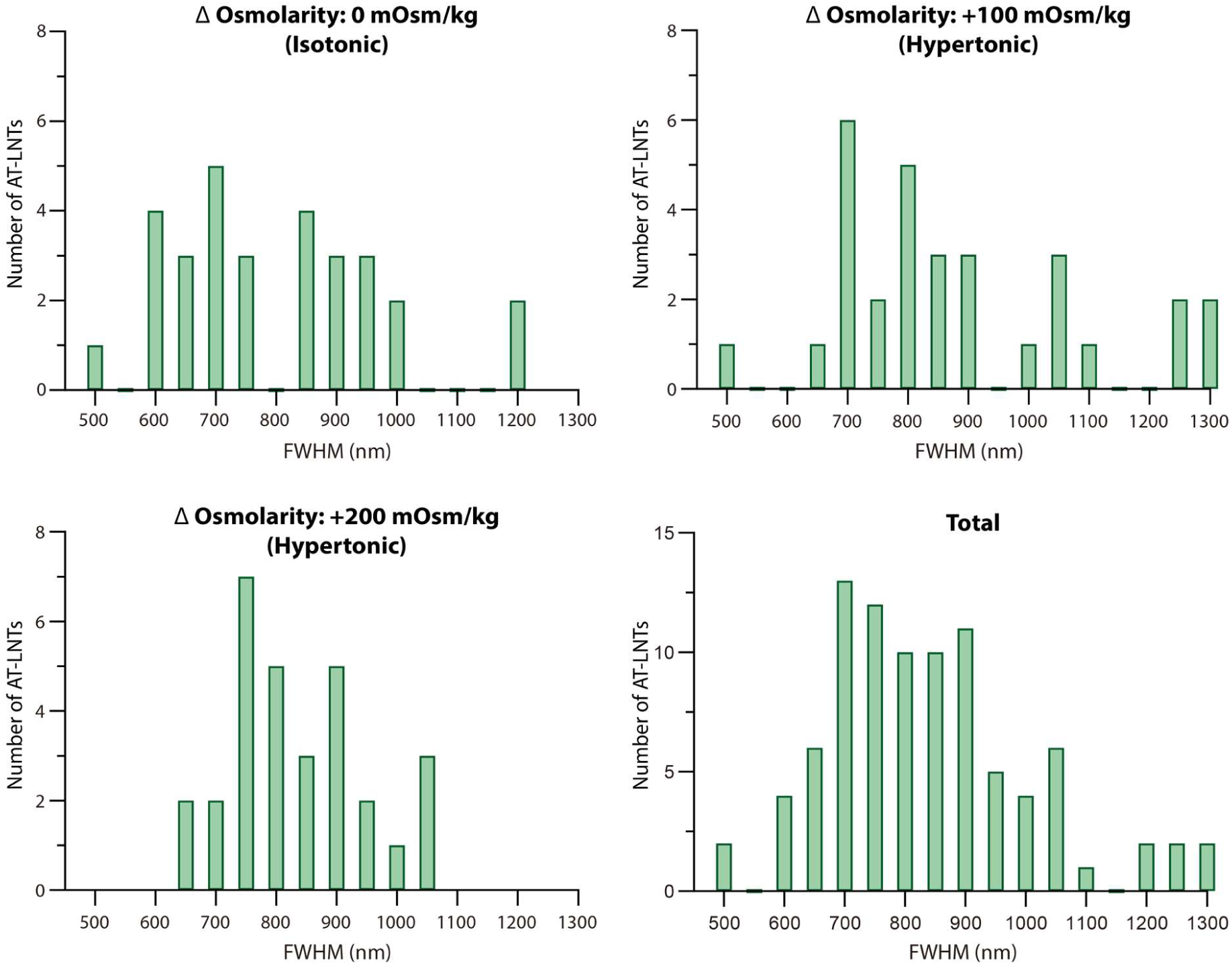
FWHM distributions of AT-LNTs. Cross-sectional FWHM distributions of lipid signals from AT-LNTs under osmotic conditions of 0 (isotonic), +100 (hypertonic), +200 (hypertonic) mOsm/kg, and the combined distribution (bottom right). Data are binned at 50 nm intervals. n = 30 measurements per condition.

**Fig. S15.**
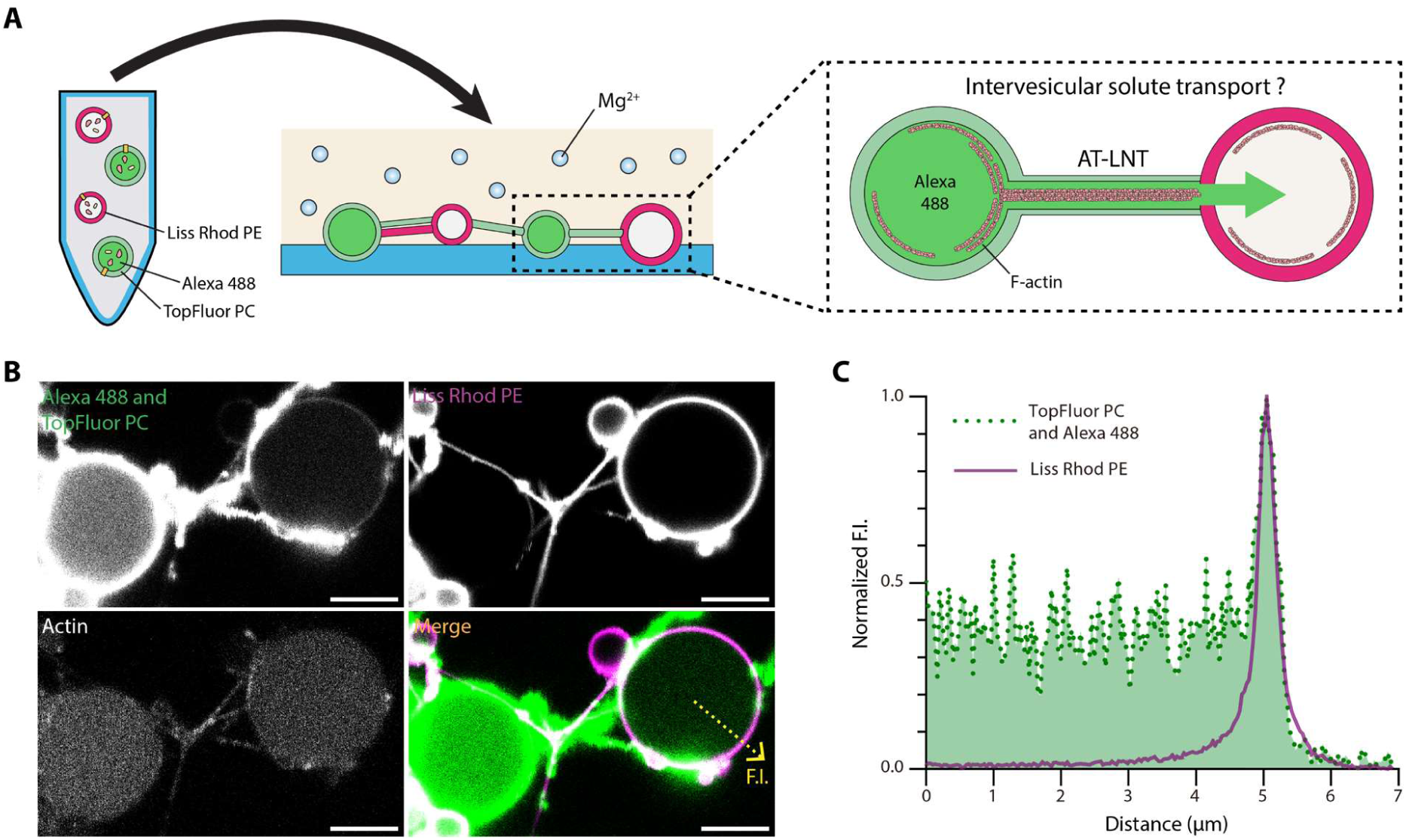
Intervesicular transfer of Alexa 488 via AT-LNT connection. **(A)** Schematic of the experimental design. TopFluor^TM^ PC-labeled (green) actin-vesicles encapsulating Alexa 488 (green) and 18:1 Liss Rhod PE-labeled (magenta) actin-vesicles without Alexa 488 were homogenously mixed and introduced into 20 mM Mg^2+^ buffer to form AT-LNT connections. **(B)** Representative confocal fluorescence images showing the presence of Alexa 488 fluorescence in the magenta vesicle connected to a green vesicle via AT-LNTs. Actin signals (bottom left) colocalize along the AT-LNT membrane visible in both green and magenta channels. The bottom right panel shows a merged image of the green (top left) and magenta (top right) channels. Scale bars, 5 μm. **(C)** Normalized fluorescence intensity profiles of the green and magenta channels measured along the yellow dashed arrow in (B). The green signal was detected in the lumenal region of the magenta vesicle, indicating intervesicular transfer of Alexa 488.

**Fig. S16.**
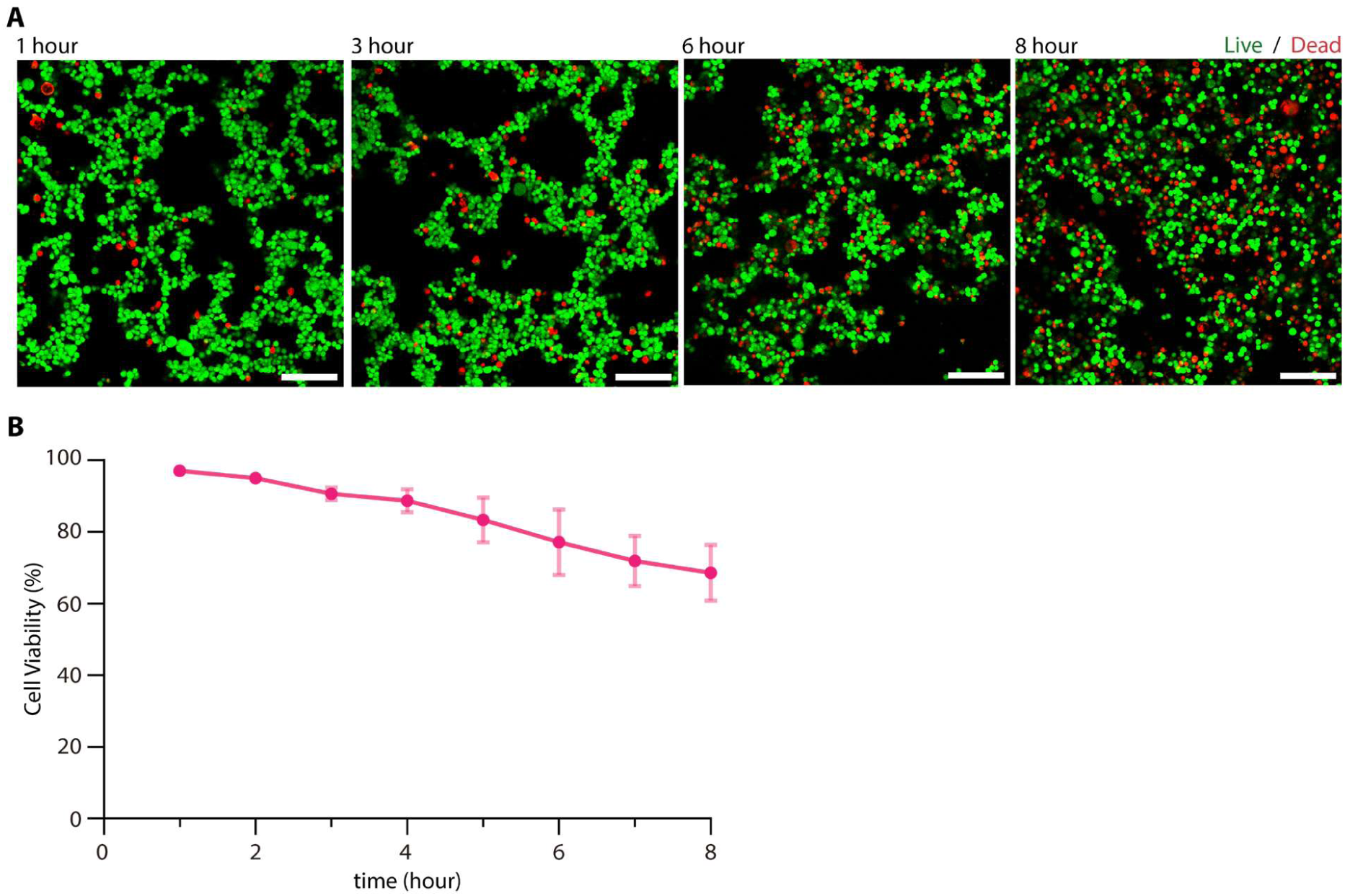
Cell viability in 20 mM Mg^2+^ buffer. HEK293 cells stained with a live/dead cell viability staining kit (green, Calcein AM; red, BOBO-3 Iodide) were introduced into the standard 20 mM Mg^2+^ external buffer. Green cells represent live cells and red cells represent dead cells. **(A)** Representative confocal fluorescence images of HEK293 live/dead cells at 1, 3, 6, and 8 h after Mg^2+^ exposure. Scale bars, 100 μm. **(B)** Cell viability over time, measured by the proportion of live (green) cells out of the total cell count. More than 60% of cells remained viable for up to 8 h. Data represent mean ± SD; n = 3 samples (581.25 μm × 581.25 μm each) for each time point.

**Movie S1.**

**Light-induced thermal undulation and retraction of intervesicular lipid nanotubes.** Intervesicular LNTs formed from actin-vesicles in 20 mM Mg^2+^ buffer were illuminated with a strong FITC lamp (at t = 0) to excite fluorescence of the membrane lipid dye (green) at the corresponding excitation wavelength. Upon illumination, LNTs undergo thermal undulations, and some intervesicular LNT connections subsequently break and retract back into the vesicles. Scale bar, 10 μm.

**Movie S2.**

**Real-time formation of an intervesicular LNT connection.** A single LNT connection forms (at t = 3 min 52 sec) from actin-vesicles following exposure to 20 mM Mg^2+^ buffer (at t = 0). Actin bundles are observed both within vesicles and along the LNTs. Green, lipid; red, actin. Scale bar, 5 μm. Representative time-lapse images are also shown in **Fig. 1B**.

**Movie S3.**

**Formation of an AT-LNT branch through inosculation.** Upon exposure of actin-vesicles to 20 mM Mg^2+^ buffer (at t = 0), a branched AT-LNT connection formed as a growing AT-LNT (visible at t = 5 min 9 sec) inosculated with a preexisting AT-LNT (t = 6 min 32 sec). The branched AT-LNT connection was subsequently pulled toward the inosculating LNT (t = 36 min 44 sec). Green, lipid; red, actin. Scale bar, 5 μm. Representative time-lapse images are also shown in **fig. S4**.

**Movie S4.**

**Multiple AT-LNTs triggered at growth/docking sites.** Upon exposure of actin-vesicles to 20 mM Mg^2+^ buffer (at t = 0), 1^st^ AT-LNT connection formed (at t = 3 min 53 sec), 2^nd^ AT-LNTs developed (t = 4 min 55 sec), an additional 3^rd^ AT-LNT develop from the growth site of the 2^nd^ AT-LNT (at t = 5 min 26 sec), and 2^nd^ and 3^rd^ AT-LNTs subsequently elongated to dock onto the previously connected vesicle. Green, lipid; red, actin. Scale bar, 10 μm. Representative time-lapse images are also shown in **fig. S5A**.

**Movie S5.**

**Bundling of an AT-LNT connection by triggered development at a docking site.** An initial AT-LNT developed and docked onto a neighboring actin-vesicle (t = 41 sec), a new AT-LNT subsequently developed (t = 7 min 33 sec), elongated (t = 9 min 26 sec), bundled with the initial AT-LNT (t = 14 min 14 sec to 18 min 42 sec), and docked onto the counterpart actin-vesicle (t = 27 min 7 sec). Time indicates duration after imaging began. Green, lipid; red, actin. Scale bar, 10 μm. Representative time-lapse images are also shown in **fig. S5B**.

**Movie S6.**

**Branching of additional AT-LNT developed from a growth/docking site.** An additional AT-LNT developed and elongated (t = 12 min 16 sec), docked onto a neighboring vesicle (t = 17 min 35 sec), detached (t = 17 min 56 sec), and inosculated with an preexisting AT-LNT (t = 23 min 46 sec) to form a branched configuration. Time indicates duration after Mg^2+^ exposure. Green, lipid; red, actin. Scale bar, 10 μm. Representative time-lapse images are also shown in **fig. S5C**.

**Movie S7.**

**Formation of intervesicular AT-LNT connections via vesicle escape.** A smaller vesicle was preformed inside a larger vesicle. The internal vesicle attached to the inner leaflet of the larger vesicle (t = 3 min 36 sec); an AT-LNT then developed from the internal vesicle and formed connections with a preexisting AT-LNT outside (t = 4 min 17 sec); and the internal vesicle subsequently escaped from the larger vesicle (t = 9 min 05 sec). Time indicates duration after imaging began. Green, lipid; red, actin. Scale bar, 10 μm. Representative time-lapse images are also shown in **fig. S6**.

**Movie S8.**

**Dynamic behavior of an AT-LNT network.** An intervesicular network of AT-LNT connections undergoes dynamic rearrangement. AT-LNTs develop, elongate, bundle, branch, and converge. Vesicles and the network are pulled along AT-LNTs, resulting in dislocation of vesicles and detachment of AT-LNTs. Time indicates duration after exposure of actin-vesicles to 20 mM Mg^2+^. Green, lipid; red, actin. Scale bar, 20 μm.

**Movie S9.**

**3D reconstruction of intervesicular AT-LNT network.** A 3D reconstruction of an intervesicular AT-LNT network was generated from Z-stacks of confocal fluorescence microscopy images. The movie shows a 90-degree vertical rotation, followed by a 90-degree horizontal rotation, and a cross-sectional view from above to bottom. Green, lipid; red, actin. The view from above is also shown in **fig. S7E**.

**Movie S10.**

**Mg^2+^-triggered LNTs from DOPC vesicles without actin.** Upon exposure of DOPC vesicles to 20 mM Mg^2+^ buffer (at t = 0), thermally undulating LNTs were spontaneously generated from DOPC vesicles over up to 20 min. Some LNTs formed stable intervesicular connections, but the majority disconnected, retracted, and disappeared. Green, lipid membrane. Scale bar, 20 μm.

**Movie S11.**

**Mg-LNTs formation at 5 mM Mg^2+^.** Upon exposure of actin-vesicles to 5 mM Mg^2+^ buffer (at t = 0), thermally undulating LNTs were spontaneously generated from actin-vesicles over up to 10 min. Some LNTs formed intervesicular connections transiently, but eventually all the LNTs disconnected, retracted, and disappeared. Green, lipid membrane; red, actin. Scale bar, 20 μm.

**Movie S12.**

**Mg-LNTs and AT-LNTs formation at 100 mM Mg^2+^.** Upon exposure of actin-vesicles (1 μM actin) to 100 mM Mg^2+^ buffer (at t = 0), thermally undulating Mg-LNTs were extensively generated from actin-vesicles and thick AT-LNTs were occasionally generated. The unstable nanotubes exhibited repeated reconnections, and the actin-vesicles were pulled along the nanotubes. Green, lipid membrane; red, actin. Scale bar, 20 μm.

**Movie S13.**

**Mg-LNTs formation at high ionic strength.** Upon exposure of actin-vesicles (1 μM actin) to 20 mM Mg^2+^ buffer supplemented with 90 mM NaCl (at t = 0), thermally undulating Mg-LNTs were extensively generated from actin-vesicles and formed intervesicular connections between neighboring vesicles. Some actin-vesicles were pulled along the Mg-LNTs, resulting in vesicle aggregations. Green, lipid membrane; red, actin. Scale bar, 20 μm.

**Movie S14.**

**Intervesicular Mg-LNT formation at high ionic strength.** Mg-LNTs extending from two neighboring actin-vesicles meet (t = 20 sec) and align toward the shortest distance (t = 8 min 55 sec). The intervesicular Mg-LNT connection thickens (t = 9 min 16 sec) and remains stable. Time indicates duration after imaging began. Green, lipid; red, actin. Scale bars, 10 μm. Representative time-lapse images are also shown in **fig. S12B**.

**Movie S15.**

**Mg-LNTs formation at a low actin concentration.** Upon exposure of actin-vesicles containing 0.05 μM actin to 20 mM Mg^2+^ buffer (at t = 0), thermally undulating Mg-LNTs were extensively generated from actin-vesicles over up to 40 min. The Mg-LNTs formed intervesicular connections, but the majority disconnected, retracted, and disappeared. Green, lipid membrane; red, actin. Scale bar, 20 μm.

**Movie S16.**

**Helical unwinding of an AT-LNT bundle by photodamage.** Upon strong illumination with a FITC lamp (at t = 0) targeting a preformed intervesicular AT-LNT bundle connection, the green iAT-LNT unwounded and retracted toward its source vesicle while alternating its position with respect to the magenta iAT-LNT. Green, Topfluor^TM^ PC lipid; magenta, 18:1 LissRhod PE lipid. Scale bar, 10 μm.

**Movie S17.**

**Formation of AT-LNT connections between an actin-vesicle and a HEK293 cell.** Actin-vesicle labeled with a membrane dye (green) and a HEK293 cell labeled with intracellular actin (red) were homogenously mixed and then exposed to 20 mM Mg^2+^ buffer at t = 0. An AT-LNT elongated from the actin-vesicle and docked onto the HEK293 cell to form a vesicle–cell connection (t = 24 min 34 sec). Soon after, a second AT-LNT connection formed (t = 25 min 15 sec). The cell interacted with AT-LNTs via a pulling motion while intracellular actin was recruited to the docking sites (t = 26 min – 59 min). One of the connections detached (t = 59 min 25 sec). Scale bar, 10 μm. Representative time-lapse images are also shown in **Fig. 6B**.

**Movie S18.**

**Elongated AT-LNT connections between an actin-vesicle and a HEK293 cell.** Actin-vesicles labeled with a membrane dye (green) and actin (red) were mixed with HEK293 cells and then exposed to 20 mM Mg^2+^ buffer. A long-distance AT-LNT connection between an actin-vesicle and HEK293 cells was observed after overnight incubation and remained intact for 4 h of observation. Scale bars, 50 μm.

**Data S1. Tabulated data for Fig. 1D, E, and G**.

**Fig. 1D**. Normalized fluorescence intensity profiles of the lipid and actin signals tabulated as a function of distance (µm) across AT-LNTs.

**Fig. 1E**. FWHM values (nm) of Gaussian fits to lipid and actin fluorescence intensity profiles measured across individual AT-LNTs (n = 30 nanotubes). Mean ± SD and *p*-value (two-tailed *t*-test) are reported.

**Fig. 1G**. Thickness (nm) of LNTs and their internal actin bundles of each AT-LNT measured by cryo-EM (n = 12 nanotubes). *p*-value is reported for the paired comparison.

**Data S2. Tabulated data for Fig. 2, C, G, and H**.

**Fig. 2C**. Normalized fluorescence intensity profiles of the lipid and actin signals tabulated as a function of distance (µm) across the dotted arrow in Fig. 2C, from center of L_o_ PC vesicle to external area.

**Fig. 2G**. FWHM values (nm) of Gaussian fits to lipid and actin fluorescence intensity profiles measured across individual AT-LNTs under varying osmolarity differences (−200, −100, 0, +100, and +200 mOsm/kg; n = 10, 10, 30, 30, and 30 nanotubes, respectively). Mean ± SD and *p*-values (two-tailed *t*-test) are reported.

**Fig. 2H**. Number of AT-LNTs per vesicle under varying osmolarity differences (−200, −100, 0, +100, and +200 mOsm/kg; *n* = 3 independent experiments per condition).

**Data S3. Tabulated data for Fig. 3, B and E**.

**Fig. 3B**. FWHM values (nm) of Gaussian fits to lipid and actin fluorescence intensity profiles measured across individual AT-LNTs at varying ionic strengths (150, 105, 82.5, and 60 mM; *n* = 10 nanotubes per condition). NA indicates that actin signal was not detected (Mg-LNT regime). Mean ± SD are reported.

**Fig. 3E**. FWHM values (nm) of Gaussian fits to lipid and actin fluorescence intensity profiles measured across individual AT-LNTs at varying actin concentrations (0, 0.01, 0.05, 0.1, 0.5, 1, and 2.5 µM; *n* = 10 nanotubes per condition). NA indicates that actin signal was not detected. Mean ± SD and *p*-value (two-tailed *t*-test, lipid FWHM between 0.05 and 0.1 µM actin) are reported.

**Data S4. Tabulated data for Fig. 4D**.

**Fig. 4D**. Normalized fluorescence intensity (FI) profiles of Liss Rhod PE and TopFluor^TM^ PC signals tabulated as a function of distance along an AT-LNT bundle and relative position (µm) with respect to Liss Rhod PE peak, measured at 12 successive positions along the nanotube axis by 1 µm gaps.

**Data S5. Tabulated data for Fig. 5C**.

**Fig. 5C**. Normalized fluorescence and bright-field (BF) intensity profiles of Alexa 488 and BF signals tabulated as a function of distance (µm) across an AT-LNT, corresponding to the line profile indicated by the dashed arrow in the ROI inset of Fig. 5C.

**Data S6. Tabulated data for fig. S1C.**

**Fig. S1C.** Fluorescence intensity of Mg²⁺ indicator as a function of time (sec) measured inside actin-vesicles under three conditions: 20 mM external Mg²⁺ with ionophore [(+) iono], 20 mM external Mg²⁺ without ionophore [(−) iono], and 0 mM external Mg²⁺ with ionophore [(+) iono] (*n* = 5, respectively). NA indicates the leakage of Mg^2+^ indicator by vesicle adhesion to substrate.

**Data S7. Tabulated data for fig. S2, B, C, and G**.

**Fig. S2B.** Normalized fluorescence intensity (FI) profiles of the PC lipid and actin signals tabulated as a function of distance (µm) across an actin-vesicle of L_d_ PC membrane, corresponding to the line profile indicated by the dotted arrow in fig. S2A.

**Fig. S2C.** Normalized FI of lumenal and cortical actin signals tabulated as a function of time (sec) after Mg^2+^ exposure of PC lipid vesicles (*n* = 5, respectively). Mean ± SD are shown.

**Fig. S2G.** Fluorescence intensity (A.U.) profiles of actin, L_o_ PC lipid, and L_d_ PG lipid signals tabulated as a function of distance (µm) across a phase-separated actin-vesicle, measured across (i) the L_o_ PC domain and (ii) the L_d_ PG domain, corresponding to the dotted arrows indicated in fig. S2F.

**Data S8. Tabulated data for fig. S7, B, C, and D**.

**Fig. S7B.** Normalized fluorescence intensity (FI) profiles of the lipid and actin signals tabulated as a function of distance (µm) across an AT-LNT at three positions (i, ii, and iii) along the nanotube, corresponding to the dotted lines indicated in fig. S7A.

**Fig. S7C.** FWHM values (µm) of Gaussian fits to lipid and actin fluorescence intensity profiles measured across individual AT-LNTs at successive positions along the nanotube axis (*n* = 11 positions). Mean ± SD and relative SD (%) are reported.

**Fig. S7D.** Pearson correlation coefficient (*r* = 0.696, 95% CI: 0.165–0.914, *R*² = 0.484) between lipid and actin FWHM values along AT-LNTs. *P* = 0.017 (two-tailed).

**Data S9. Tabulated data for fig. S11, B and C.**

**Fig. S11B.** FWHM values (nm) of Gaussian fits to lipid fluorescence intensity profiles measured across individual Mg-LNTs and AT-LNTs at 5 mM and 10 mM Mg²⁺, respectively (*n* = 10 nanotubes per condition). Mean ± SD are reported.

**Fig. S11C.** Proportions (%) of Mg-LNTs and AT-LNTs formed at 5, 20, and 100 mM Mg²⁺ (*n* = 3 independent experiments per condition).

**Data S10. Tabulated data for fig. S14.**

**Fig. S14.** Frequency distributions of lipid FWHM values (nm) of AT-LNTs under isotonic conditions, hypertonic conditions (+100 and +200 mOsm/kg), and pooled across all conditions, binned at 50 nm intervals.

**Data S11. Tabulated data for fig. S15C.**

**Fig. S15C.** Normalized fluorescence intensity (FI) profiles of Liss Rhod PE and TopFluor^TM^ PC/Alexa 488 signals tabulated as a function of distance (µm) from center to external space of a Liss Rhod PE-labeled actin-vesicle, corresponding to the dotted arrow in fig. S15B.

**Data S12. Tabulated data for fig. S16B.**

**Fig. S16B.** Cell viability (%) of HEK293 cells tabulated as a function of time (hours) following 20 mM Mg²⁺ treatment, assessed by live/dead staining (*n* = 3 independent experiments). Mean ± SD are shown.

